# Inferring post-transcriptional regulation within and across cell types in human testis

**DOI:** 10.1101/2024.10.08.617313

**Authors:** Saad Khan, Megan Elcheikhali, Andrew Leduc, R Gray Huffman, Jason Derks, Alexander Franks, Nikolai Slavov

## Abstract

Single-cell tissue atlases commonly use RNA abundances as surrogates for protein abundances. Yet, protein abundance also depends on the regulation of protein synthesis and degradation rates. To estimate the contributions of such post transcriptional regulation, we quantified the proteomes of 5,883 single cells from human testis using 3 distinct mass spectrometry methods (SCoPE2, pSCoPE, and plexDIA). To distinguish between biological and technical factors contributing to differences between protein and RNA levels, we developed *BayesPG*, a Bayesian model of transcript and protein abundance that systematically accounts for technical variation and infers biological differences. We use *BayesPG* to jointly model RNA and protein data collected from 29,709 single cells across different methods and datasets. *BayesPG* estimated consensus mRNA and protein levels for 3,861 gene products and quantified the relative protein-to-mRNA ratio (rPTR) for each gene across six distinct cell types in samples from human testis. About 28% of the gene products exhibited significant differences at protein and RNA levels and contributed to about 1, 500 significant GO groups. We observe that specialized and context specific functions, such as those related to spermatogenesis are regulated after transcription. Among hundreds of detected post translationally modified peptides, many show significant abundance differences across cell types. Furthermore, some phosphorylated peptides covary with kinases in a cell-type dependent manner, suggesting cell-type specific regulation. Our results demonstrate the potential of inferring protein regulation in from single-cell proteogenomic data and provide a generalizable model, *BayesPG*, for performing such analyses.

## Introduction

Understanding the mechanisms that establish the identity of individual cells from complex tissues is a fundamental biological question. These mechanisms include both transcriptional and post-transcriptional regulatory processes. Transcript abundance and regulation have been extensively studied over the last decade, including both cell atlases^1,2^ and inference of transcriptional regulation^3,4^. In contrast, the exploration of post-transcriptional regulation via protein synthesis and degradation has been limited by the challenges of reliably and scalably quantifying proteins in single cells^5–9^. This limitation is increasingly mitigated by single-cell mass spectrometry (MS) methods that support multiplexing^10^, thus enabling quantitative protein analysis across thousands of single human cells^11,12^. These methods can sample millions of peptide copies per cell^13,14^ across many single cells, which can support the quantitative exploration of post-transcriptional regulation as reflected in the differences between protein and RNA levels. These differences likely stem from regulation of protein synthesis and degradation^15–17^.

Such data interpretation also necessitates approaches for jointly modeling mRNA and protein levels while accounting for different sources of variability and thus distinguishing biological regulation from technical noise^18,19^. To accomplish this goal, we developed a Bayesian ProteoGenomic (*BayesPG*) framework for modeling single-cell RNA-seq and MS data at the level of cell types. *BayesPG* models abundance at the level of cell types to help with the coarser problem of understanding variation across cell types. By modeling the technical variability (within and across data sets) and biological variability (across modalities and cell types), *BayesPG* can systematically identify proteins whose abundance cannot be explained solely by the abundance of their corresponding mRNAs. Such identification requires that the data provide sufficient statistical powers: post-transcriptional regulation cannot be quantified when the technical variation in mRNA or protein estimates is significantly larger than the variation due to underlying biological processes.

We apply *BayesPG* to explore post transcriptional regulation in single cells from human testis. Previous work has shown that, alongside brain tissues, the testis show the highest disagreement between protein and mRNA levels^20^. We focus our work on the testis, not only due to this disagreement but also because the disagreement is driven, at least in part, by the complex and highly specialized processes undertaken by the cells in the testis (eg; maturation of sperm cells from spermatogonial stem cells). Analysis is performed on 6 cell types, including endothelial cells (EC), peritubular myoid cells (PTM), leydig cells (LC), spermatagonia cells (SPG), spermatocytes (SPC) and spermatids (St).

## Results

### Data sets and alignment

Observed differences in mRNA and protein levels are driven by both biological and technical factors. To compensate for the technical factors and isolate the biological ones, we need measurements collected in different ways, ideally with near orthogonal biases^18,21^. To this end, we acquired proteomics data with very different methods to analyze about 6,000 single cells and compiled published single-cell mRNA-seq data^22,23^ from about 24,000 single cells, Fig. 1. These RNA data were collected using similar technologies (10x Genomics and DropSeq) but were generated across multiple donors, sequenced at different depths (across and within study), contained technical replicates and cell types were annotated by different teams and markers.

**Figure 1.**
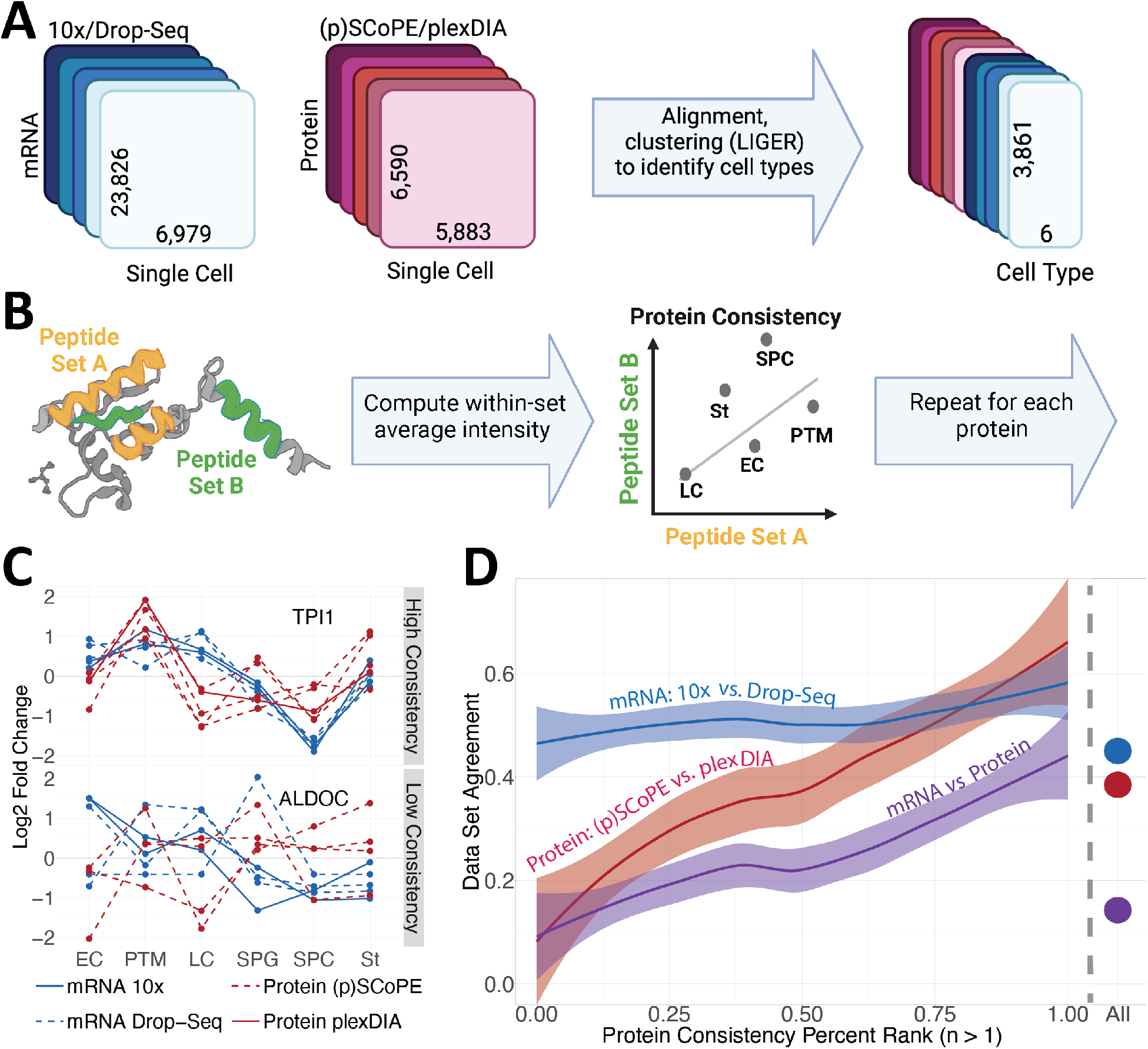
Dataset alignment and reliability assessment. **A** Overview of model preparation. The input data contain multiple datasets with observed transcript abundances and peptide-level intensities for 12,862 single cells. LIGER is used to perform alignment and clustering into cell types. Cell type labels are retained and untransformed transcript abundances and peptide intensities are summarized across cells assigned to each type. **B** Peptides originating from the same proteins are used to estimate the consistency of protein quantification for proteins with two or more peptides quantified. We randomly sort quantified peptides into two sets, and the cluster average abundance for the two sets are correlated across cell types to obtain consistency estimates. **C** Examples of proteins measured with high and low protein consistency, displayed in top and bottom panels, respectively. These proteins also have high and low data set agreement, respectively, depicted by fold changes observed for each dataset. **D Left** Across cluster correlation between data sets as a function of the protein consistency estimated as outlined in panel B. **D Right** Data set agreement is averaged across all gene products; The agreement for proteins for which only one peptide is consistently quantified (and thus have not consistency estimates) are shown in the right panel labeled with All.

Our single-cell proteomic data sets were collected analysing single-cell suspensions with 3 different methods. The dissociated tissue was collected from a single human donor, and thus the data does not capture variability across human subjects, though it is smaller than the variability between cell clusters, Extended Data Fig. 1. We acquired 5 datasets. Four datasets used isobaric mass tags and carriers and perform quantification at the MS2 level; one of these datasets used prioritized data acquisition that selects peptides using a tiered prioritization list^24^. Our fifth data set was acquired using plexDIA, where quantification is carried out at the MS1 level and no carrier is used^25,26^. Since these methods perform quantification based on different ions (precursors vs. reporter ions) quantified at different stages (MS1 vs. MS2) their biases are distinct^27^. Thus, their measurements allow distinguishing between biological variance (shared across methods and datasets) and technical variance (specific to methods and datasets).

To jointly analyze the protein and RNA data, we started by aligning the datasets to generate cross modality clusters, Fig. 2A. The cells within each cluster are assigned a cell type based on a consensus of the independent mRNA annotations, Extended Data Fig. 2B. Aligning mRNA and protein data is challenging, as mRNA and protein levels in cells from the same type may systematically differ due to both technical and biological factors. To mitigate these challenges, we chose a feature space for the alignment that contains gene products exhibiting similar covariation within each modality, Extended Data Fig. 2A. Within this feature space, we use non-negative matrix factorization for alignment, as implemented in LIGER^28^, because it explicitly factors shared and not shared sources of variation across data sets.

**Figure 2.**
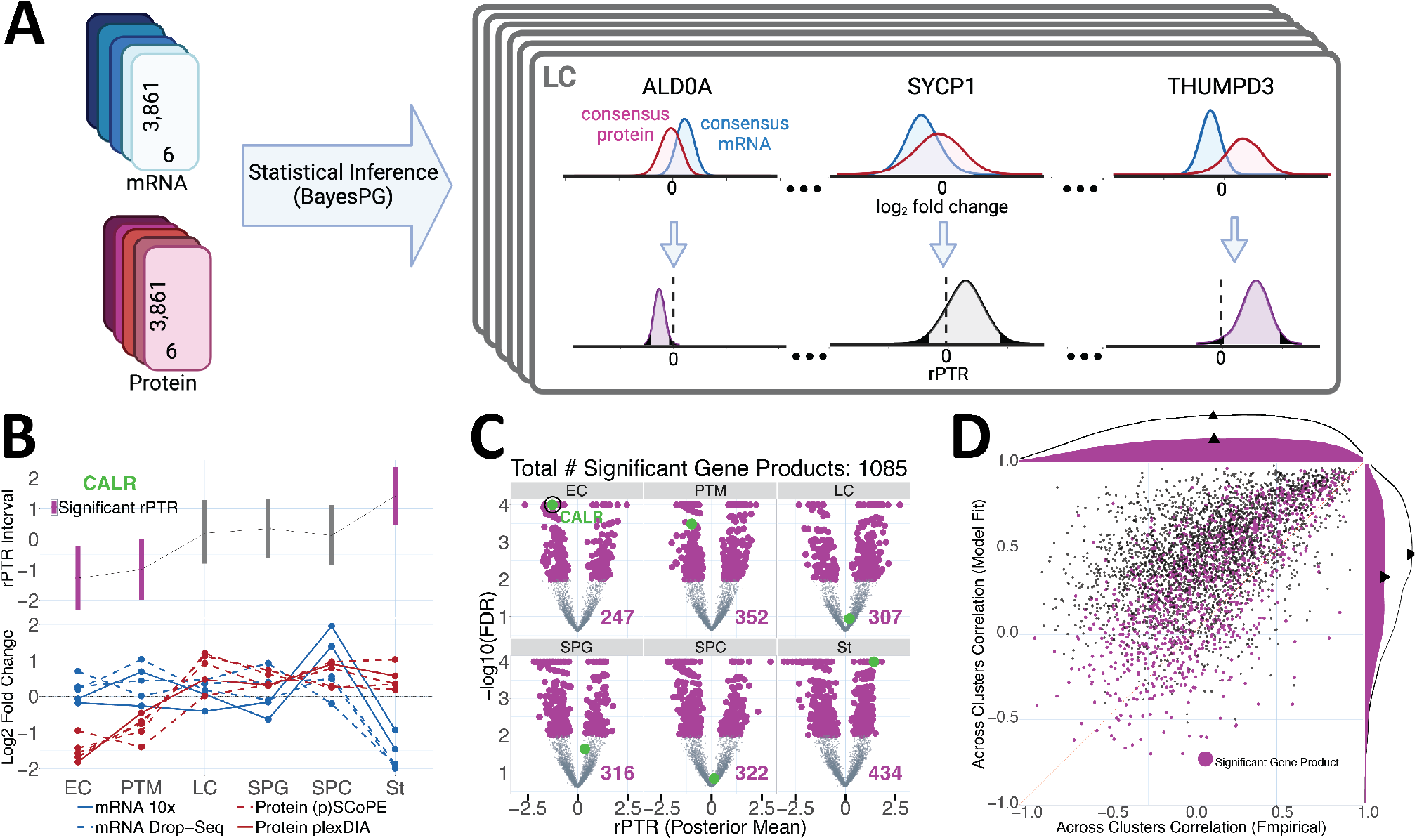
Global quantification of cell-type specific post-transactional regulation of individual proteins. **A** Examples of consensus posterior distributions inferred by *BayesPG* for the protein and RNA products of 3 genes in the Leydig cell (LC) cluster. The ability to detect significant rPTR (in purple) for a given protein depends on the width of posterior intervals corresponding to consensus mRNA and protein. This width is inversely proportional with the agreement across peptides associated with each protein and across data sets within each modality. For example, nonzero rPTR can be detected for ALDOA as posterior intervals are narrow. Meanwhile, the wide marginal posterior distributions associated with SYCP1 indicate low confidence and no significant rPTR. **B** An example comparison of rPTR posterior interval and observed *z*-transformed averages for Calreticulin RNA and protein. The top panel displays the posterior interval for rPTR for each cell type, with purple lines representing significant cell types. The bottom panel displays *z*-transformed cluster-level averages of log_2_ observed values. Corresponding *z*-transformations are applied separately for each data set and gene product across cell types. **C** Posterior means of model fit rPTR displayed on x-axis, and − log_10_ posterior expected proportion of false discoveries for gene-level testing of posterior samples of rPTR against across cell types mean. Purple points represent genes with significant nonzero rPTR relative to across cell types average. The green point in each panel corresponds to Calreticulin. **D** Comparison of model-fit and empirical across clusters mRNA, protein correlations among gene products observed in three or more clusters for both modalities. Purple points denote gene products with significant rPTR in at least one cell type. The purple marginal density plots reflect posterior mean correlations across significant gene products, while correlations across all gene products (including those with significant rPTR) are shown with the black marginal density lines. Correlations of posterior mean values are consistently higher than empirical correlations. This indicates that *BayesPG* resolves attenuation caused by noise and technical variability, and avoids the presumption of post transcriptional regulation by biasing rPTR toward 0.

Since cell-type clusters are inferred using our alignment procedure, we first qualitatively confirm the clustering by cell type in the space of PC1 and PC2 for the proteomic data, Extended Data Fig. 2. Next, we compare gene product correlations across cell types in the alignment space to quantify the agreement of relative levels across modalities, Extended Data Fig. 2B. We quantitatively evaluate cluster compactness for the proteomic data by comparing the within to between cluster ratio of Mahalanobis distances (Extended Data Fig. 2B). With cell type labels assigned to cells in all data sets, we then created gene by cell type data sets by aggregating over cells associated with each cluster, Fig. 1A. These results support the overall quality of the alignment, Extended Data Fig. 2.

### Characterizing measurement reliability

Given abundance measures in six cell-type clusters for each mRNA and protein data set (Fig. 1A), we then turn to understanding the drivers underlying apparent differences between these data sets. In particular, accurate characterization of post transcriptional regulation first requires an understanding of the degree to which different technical factors influence variability of observed quantification^11,18^. One way to measure the impact of technical factors is by computing measurement reliability^21^. To estimate the reliability associated with each gene product, we compare two reliability estimates: (i) protein consistency and (ii) data set agreement. We define protein consistency as the agreement between different peptides originating from the same protein within a data set, Fig. 1B. In contrast, data set agreement reflects the consistency of quantification between different measurement techniques. Reliability estimation by these measures is performed at the cluster level by correlating transcript count sums for mRNA and, for protein data, the relative log_2_ peptide intensity across cells associated with each cluster.

Both protein consistency and dataset agreement are computed by correlating log fold change measurements across clusters. To compute protein consistency, we randomly split peptides associated with each protein into two groups, compute the average within each group, and correlate across cell types, Fig. 1B. Protein consistency is a useful metric for data collected via bottom-up proteomics since measurements of multiple peptides provide mostly independent estimates for the abundance of the protein from which they originate^11^. While measuring protein consistency provides insight into technical factors affecting protein quantification, it is only possible to compute this statistic for proteins with two or more peptides observed and not applicable to other modalities. In contrast, dataset agreement, estimated as the across cluster correlation of log fold changes from different datasets, can be computed with measurements of any type, including protein and RNA abundance data, and between the two modalities (“across-modality agreement”). Dataset agreement between (p)SCoPE and plexDIA is particularly informative because protein abundance is measured with near orthogonal technical biases, as peptide quantification is performed on very different ions (reporter and precursor ions for (p)SCoPE and plexDIA, respectively)^11,19^. In contrast, biases in measuring relative RNA abundance that are shared between 10x and DropSeq will result in overestimating RNA reliability.

Next, we compared our two estimates of reliability, protein consistency and dataset agreement. A detailed comparison for TPI1 and ALDOC indicates that the consistency of quantification from different peptides corresponds to the level of dataset agreement, Fig. 1C. To test the generality of this observation, we displayed in Fig. 1D the agreement across datasets as a function of the percent rank sorted protein consistency for all proteins with multiple quantified peptides. The two protein reliability estimates strongly correlate, thus generalizing the observation from Fig. 1C and bolstering each other. The dataset agreement for proteins represented by one quantified peptide and their corresponding transcripts are shown in the right panel of Fig. 1C. The dataset agreement among transcript measurements is generally higher than for the protein datasets, which may reflect overestimation due to shared biases between the scRNA-seq methods.

Crucially, the concordance between consistency and dataset agreement suggest that our estimates capture the influence of technical factors and can support biological inferences for or the majority of proteins; For proteins with the lowest 5% consistency, protein dataset agreement is comparable to the across-modality agreement, which means that for these genes the differences between the relative mRNA and protein levels could be driven entirely by technical factors. For the majority of proteins, however, the across-modality agreements are lower than the within-modality agreements. This difference strongly indicates that most protein abundances are regulated via cell-type specific modulation of protein synthesis and degradation: If differences between modalities were solely explained by technical variation (null model), the mRNA-protein agreement would be between the 10x-DropSeq and (p)SCoPE-plexDIA dataset agreement values.

While the results in Fig. 1D support widespread post-transcriptional regulation, its reliable inference requires a dedicated approach. In particular, gene-level quantification of post-transcriptional effects necessitates novel statistical methodology, beyond simple reliability calculations, which account for technical variation both within- and between-measurements, and yields estimates of protein-to-RNA ratios. This necessity motivated the development of *BayesPG*, described below.

### Quantifying post-transcriptional regulation with *BayesPG*

To precisely and rigorously quantify biological differences between the RNA and protein products of each gene in each cell type, we need a statistical model that systematically accounts for distinct sources of variability. To this end, we propose *BayesPG*, a model for inferring the log_2_ relative protein-to-mRNA ratio (rPTR) across cell types. To infer rPTR, we use the model structure in Extended Data Fig. 3 to represent peptide levels and transcript counts for each cluster across multiple mRNA and protein datasets. To jointly infer consensus protein and mRNA levels, *BayesPG* models in log space the protein abundance as the corresponding RNA abundance plus protein-to-mRNA ratio for each gene product and cell type. We include gene-level average parameters for each gene product and focus our model on the fold changes of mRNA and protein relative to their across cluster averages. Cluster-level parameters account for systematic differences between cell types, for example, due to different cell sizes. *BayesPG* accounts for technical variability across peptides associated with each protein using a protein and dataset level sampling variance parameter. For transcript data, *BayesPG* includes a gene-level overdispersion parameter to allow for increased variance of transcript counts relative to the mean. For exact model specification and technical details see Extended Data Fig. 3.

One challenge with jointly modeling RNA-seq and mass spectrometry data is that the dynamic range of transcript fold changes is typically larger than the corresponding measured dynamic range of protein fold changes. This likely occurs due to technical factors, including interference by coisolation affecting reporter ion quantification^29^ and dropouts in scRNA-seq data. To account for such dynamic range differences, for each gene product and measurement technique we include a scaling parameter; it ensures that the dynamic ranges of log_2_ fold changes across clusters for each mRNA and its corresponding protein are approximately the same. This scaling guards against mis-interpreting ratio compression artifacts as biological effects. While this scaling approach prevents false positive discoveries, it likely increases false negatives, as any post transcriptional effects that manifest predominantly as differences the between the dynamic range of mRNA and protein will not be detected. These scaling parameters and all other model parameters are visualized in Extended Data Fig. 3. We include a detailed description of the model specification and all prior distributions in Methods.

We fit our model *BayesPG* using No-U-Turn MCMC sampling^30^ and find that the model appropriately reflects essential features of observed data. To verify model fitness, we compute various posterior predictive model checks (PPCs)^31^. With PPCs, we compare statistics computed on observed data to the same statistics computed from data simulated under the proposed model; similarity between the statistics supports the validity of the model. We compare fold changes, across cluster variance and quantiles of the across cluster mRNA-protein correlations and show that *BayesPG* inferences are indeed consistent with the data along the most important dimensions, Extended Data Fig. 5. See Methods for more detail.

Having established confidence in *BayesPG*, we then characterize the consensus relative abundance and rPTR for each gene product and cell type with Monte Carlo samples as illustrated in Fig. 2A. Since rPTR is defined relative to the average across cell types, the posterior distribution for centered rPTR in a given cell type will contain zero if the effect of post transcriptional regulation cannot be distinguished from the average effect across other cell types. When a 95% posterior interval for centered rPTR excludes zero, we declare the rPTR “statistically significant” for that gene. As intuitive examples, we display the posterior distributions for the products of a few genes in Leydig cells, Fig. 2A. Some modeled gene products, e.g. ALDOA, exhibit high reliability, and thus we have the power to detect significant rPTR. In some cases, e.g. THUMPD3, the observed difference between mRNA and protein is large enough, that despite low reliability, rPTR is still inferred to be significantly different from zero. For other gene products, e.g. SYCP1, technical factors alone could account for observed mRNA-protein differences, and thus rPTR estimates are not declared statistically different from zero, Fig. 2A. This dependence between capacity to detect nonzero rPTR and data reliability at the gene product level is systematic across proteins, as demonstrated in Extended Data Fig. 4. To further exemplify *BayesPG* predictions across cell types, Fig. 2B (top) depicts the inferred 95% posterior intervals of rPTR for CALR across all cell types, along with *z*-transformed log_2_ cluster-level averages computed from the raw measurements (bottom). The data show how both the agreement across datasets and the differences in consensus RNA and protein abundances relate to identifying significant patterns of protein regulation. Applying this analysis to our entire dataset, we find that 1, 028 (28%) proteins have significant rPTR in at least one cluster, Fig. 2C.

To holistically quantify protein and RNA similarity across all cell types with increased statistical power, we estimate the correlations between protein and RNA levels across clusters for all genes, Fig. 2D. Since *BayesPG* corrects for attenuation due to measurement error, the across clusters mRNA-protein correlation (previously referred to as across-modality dataset agreement) computed using model-based consensus estimates are markedly higher than those computed from the observed data directly: the median across-modality agreement increases from 0.17 (observed) to a median of 0.49 (model), Fig. 2D. While some differences between mRNA and protein are attributed to technical effects, the model based estimates of mRNA-protein correlations indicative substantial post-transcriptional regulation, especially for RNAs with low protein correlations.

### Post-transcriptional regulation of functional protein groups

Next, we used *BayesPG* to group per-gene estimates into known functional groups defined either by the Gene Ontology (GO) or by protein complexes, Fig. 3. This grouping helps reduce posterior uncertainty and increases statistical power to detect significant post-transcriptional regulation. Specifically, we seek to identify functional groups with cell-type rPTR that is significantly different from its average rPTR across all cell types based on posterior intervals (see Methods). This approach identified 1, 453 significant GO groups and 851 complexes, and a few of them are displayed in Fig. 3A. Considering that spermatogenesis is the dynamic process during which spermatagonial stem cells differentiate into haploid spermatids, some deviations between RNA and protein may reflect delayed protein production as observed in other systems^32,33^. These effects are likely not dominant in spermatogenesis as its period of 73 days is longer than typical delays in protein accumulation.

**Figure 3.**
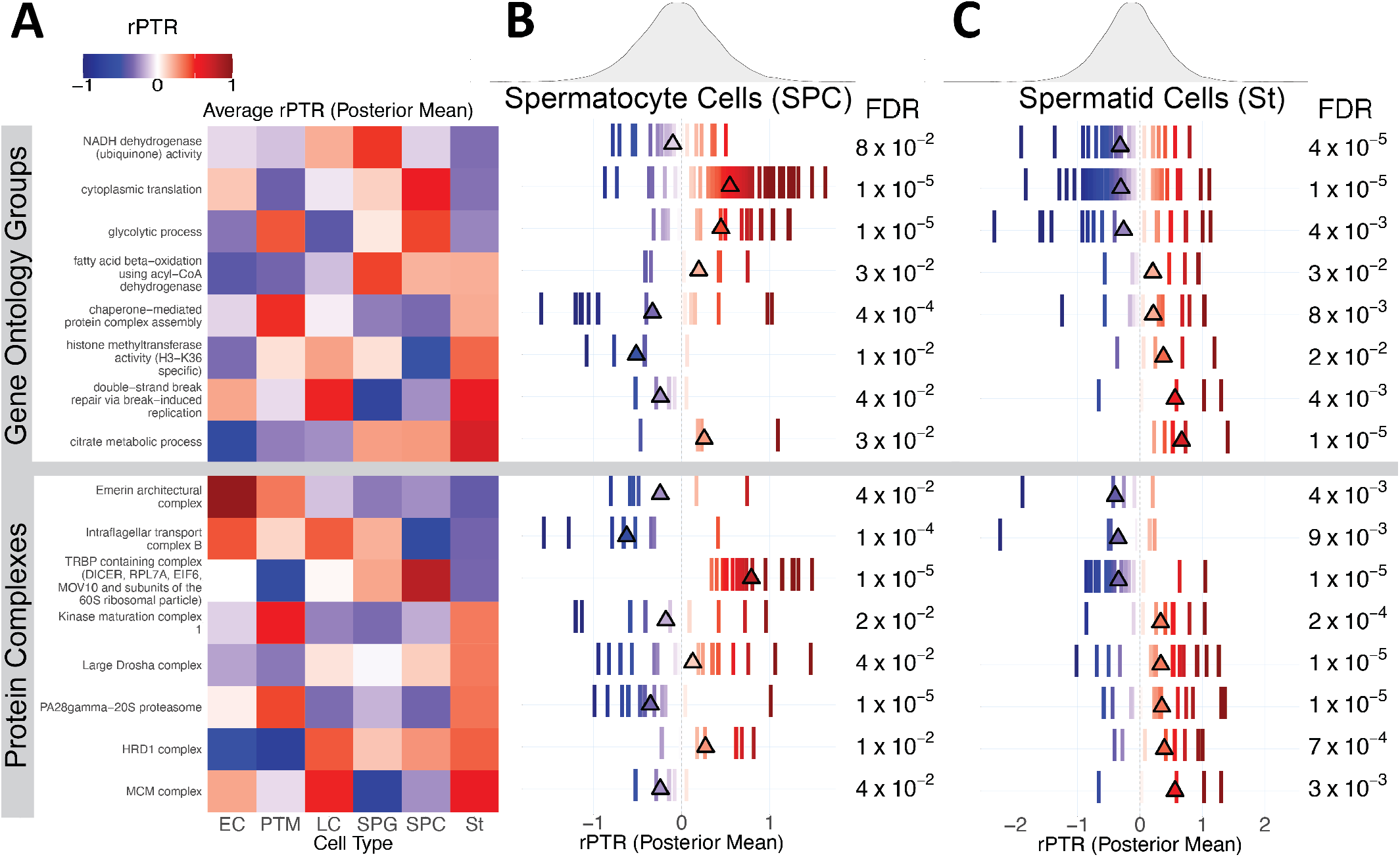
Analysis of rPTR for GO groups and protein complexes. **A** Heatmap of average posterior means of rPTR across genes associated with select significant GO groups (above line) and complexes (below line) for each cell type. Color indicates value and direction. **B** Posterior mean of rPTR averages for spermatocyte cluster among genes associated with groups and complexes displayed in heatmap. Density of group-level rPTR averages for spermatocyte cluster across all gene products is displayed in top panel. Estimated FDR corresponding to groups and complexes is shown on the left (see Methods). **C** Posterior mean of rPTR averages for spermatid cluster, as well as density across gene products and estimated FDR.

Among the proteins with the largest rPTR are those related to spermatogenesis and metabolic processes, and this suggests that protein synthesis and degradation contribute to their regulation, Fig. 3. Mitochondria are extensively remodeled during spermatogenesis^34^, and our data implicates protein synthesis or degradation in this process. Specifically, we observe that the protein to RNA ratios for mitochondrial respiratory chain complexes changes across spermatogenesis, Fig. 4. These changes have opposing trends for different cell types, Fig. 4. Such trend changes depending on data sub-setting are know as a Simpson’s Paradox and have motivated single-cell analysis^26^.

**Figure 4.**
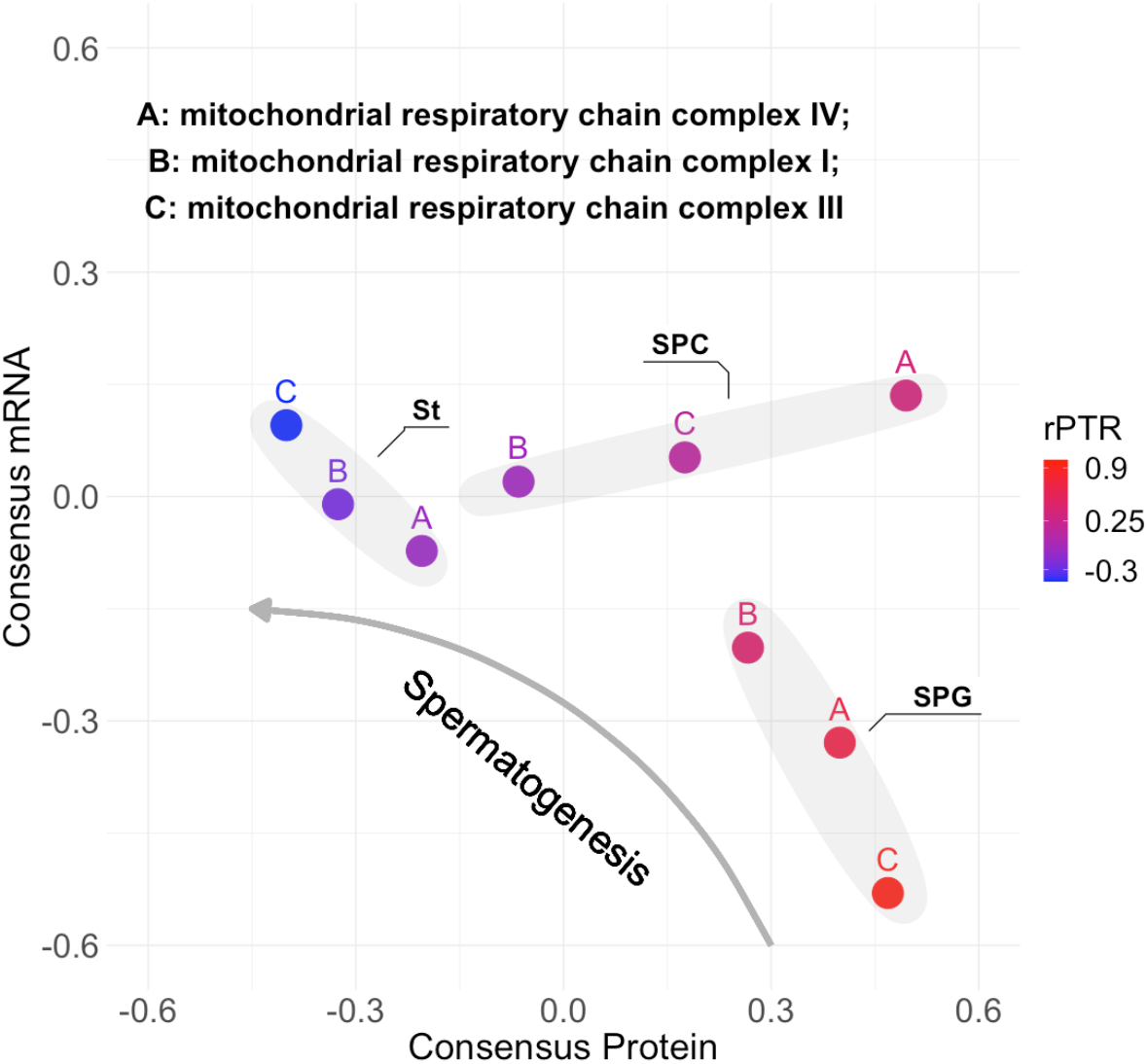
mRNA and protein variation during spermatogenesis for the mitochondrial respiratory chains. The consensus RNA and protein abundances for the indicated mitochondrial GO terms are show for 3 cells types denoted by the shaded ovals. Each data point is colored by its corresponding rPTR.

Since protein abundance decrease while mRNA abundance increases from spermatocytes to spermatids, the trends suggest increased levels of protein degradation, Fig. 4.

Other post-transcriptionally regulated processes include glycolysis and lactate metabolism. Consensus mRNA and protein levels for glycolytic enzymes change such that the spermatids maintain negative rPTR, despite having the highest consensus protein levels, Extended Data Fig. 6. This suggests that proteins involved in glycolysis are regulated after transcription during spermatogenesis. Furthermore, we find significant rPTR for ontologies relating to the previously described utilization and inter-conversion of lactate to pyruvate during spermatogenesis. We find that while consensus protein levels for lactate dehydrogenase subunit B remain lower than the mean, we see negative rPTR for spermatogonia and positive for spermatids, indicating control at the level of protein synthesis and degradation. We also note very high consensus protein levels for the testis specific lactate dehdydrogenase A-like 6B in the spermatocytes and spermatids. We observe a similar shift from somatic to germ cell specific forms for the alpha subunit of the pyruvate dehydrogenase E1 component. Interestingly, both gene products have very high protein and mRNA levels, but, are maintained at negative rPTR, implying the involvement of protein synthesis and degradation in regulating this shift to protein isoforms which are better adapted to spermatogenesis.

Our data also indicate protein to RNA changes that shift *β*-oxidation of fatty acids from the mitochondria (short-chain fatty acids) to the peroxisome (long-chain fatty acids), Fig. 3. Specifically, the rPTR decreases for *β*-oxidation of fatty acids from Spermatogonia to Spermatocytes. At the same time, the rPTR for very long chain fatty acid *β*-oxidation increases, Fig. 5C. Simultaneously, rPTR increases in peroxisome assembly factor 2 and the peroxisomal multifunctional enzyme alongside an increase in mitochondrial long chain specific acyl-coA dehydrogenase. Taken together, these results indicate enzyme level regulation mediating an overall shift towards long chain fatty acid based *β*-oxidation and a compensatory shift towards cytoplasmic beta oxidation via the peroxisomes. A related remodeling includes a significant increase in rPTR for citrate metabolic process from Spermatocytes to Spermatids. Alongside increased rPTR for citrate synthase, we find an increase in cytoplasmic Aconitase, which in high iron conditions catalyzes the rate limiting conversion of citrate to isocitrate. Concomitant to this, we observe a 2-fold shift in the rPTR for iron responsive element binding protein 2, which serves as a co-factor for cytoplasmic aconitase. These data implicate a compensatory shift towards cytoplasmic activity of critical components of the tricarboxylic acid cycle with a marked shift in the rPTR of proteins responsible for the production of citrate.

**Figure 5.**
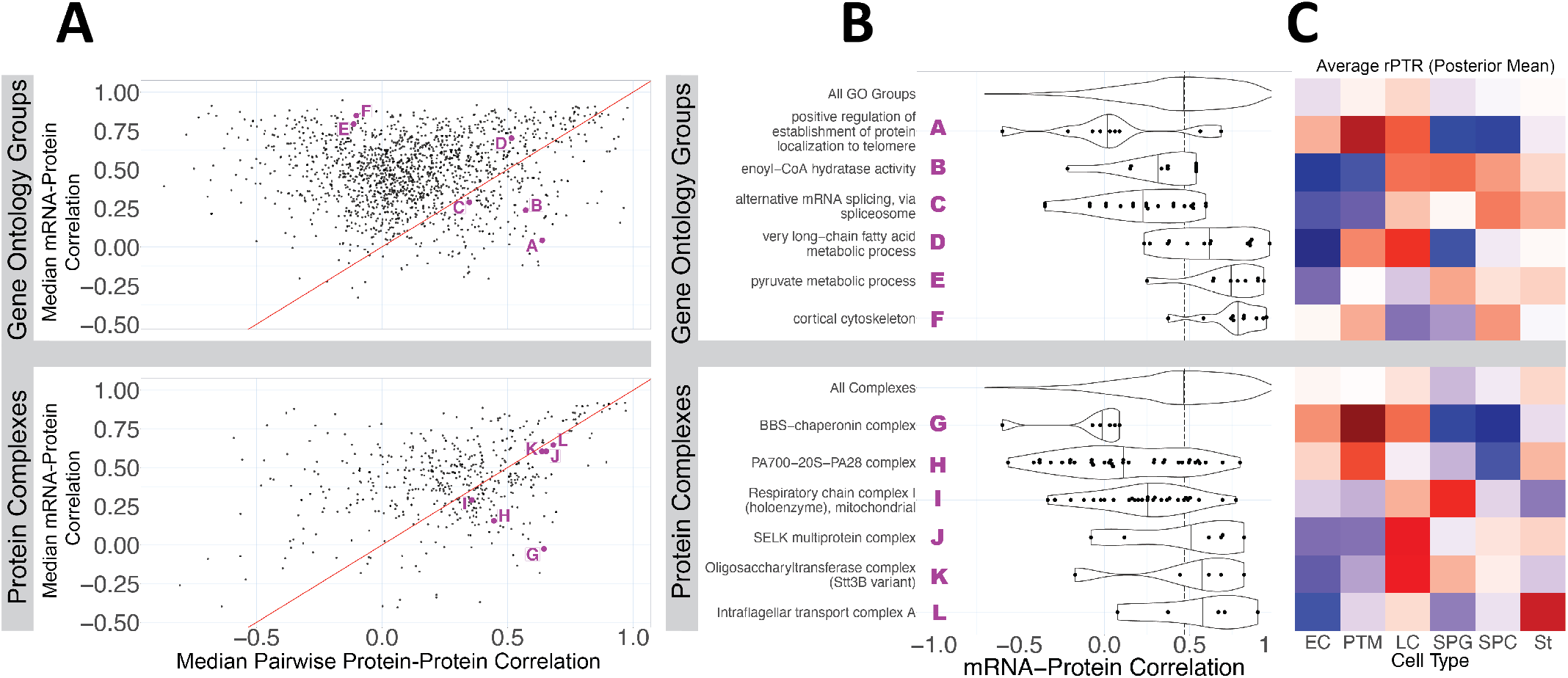
GO and Complex Analysis of Across Clusters mRNA-Protein Correlation. **A** Posterior mean of median across clusters pairwise consensus protein-protein correlation. The median is computed across pairs of gene products associated with each GO group and protein complex, displayed on the *y* − axis. One the *x* − axis we can see the median pairwise mRNA-mRNA correlation across genes products in each group. The figures on the top relate to GO groups, while the figures on the bottom relate to protein complexes. **B** Violin plot representing posterior mean of mRNA-protein correlation across clusters among gene products associated with displayed GO groups (above line) and protein complexes (below line). Points represent corresponding posterior means. **C** Heatmap of average posterior mean rPTR across gene products for associated GO groups (above line) and protein complexes (below line).

Next we focused on global trends that are well captured by RNA-protein correlations: For some proteins, the variation across cell types is strongly correlated to their corresponding mRNAs while for others it is not, Fig. 2D: Thus, we sought to test if this variation is related to the protein functions by evaluating distributions of correlations within a functional groups, Fig. 5. We observed that functions with specialized, cell type-specific roles (such as protein localization to telomeres and alternative splicing; Fig. 5B, upper panel) typically exhibited low mRNA-protein correlations, indicating greater dependence on post-transcriptional mechanisms. In contrast, GO terms central to general cellular functions (such as pyruvate metabolism and cortical cytoskeleton) displayed higher mRNA-protein correlations, suggesting a stronger influence of mRNA on protein levels. Interestingly, while these central processes showed high mRNA-protein correlation, we also observed significant shifts in the relative protein-to-mRNA ratio (rPTR) across cell types, particularly for processes such as very long-chain fatty acid and pyruvate metabolism, Fig. 5.

Next we focus on post-transcriptional regulation of subunits of protein complexes. We selected complexes whose protein subunits are strongly correlated across clusters (Fig. 5B), as typical for stable complexes. These high correlations within a complex provide additional confidence in the accuracy of our consensus protein levels. Yet, even for this subset of proteins, we observe significant deviations between consensus RNA and protein levels, including for the BBS chaperonin complex and the proteasome, Fig. 5B. Similar to our GO term analysis, we find specialized, cell type-specific complexes that show low cross-modality correlation (respiratory chain 1, Protea-some, Fig. 5B, lower panel). Within the high correlation subset, two of the three complexes can be linked to the endoplasmic reticulum (ER) and protein folding^35,36^, with Stt3B-driven N-terminal glycosylation essential for sperm maturation^36^. Similarly, we observe significant rPTR for other ER localized complexes (SELK, HRD1, Kinase Maturation). Together, these findings suggest that coordinated synthesis and degradation may contribute to the unfolded protein response in the ER. Contrasting functionally coordinated correlation of rPTR, we find the interflagellar transport complex A (IFT-A) to have high cross-modality correlation while the functionally related BBS chaperonin complex shows very low correlation (alongside very high protein-protein correlation). Within the context of our observations thus far, this suggests that the rPTR of the IFT-A is maintained across celltypes, while the BBSome shows more cell type specific deviations in ratio of protein to mRNA.

Our MS data also quantified a few hundred peptides with post-translational modifications in dataset 1, including phosphorylation, acetylation and methylation, Fig. 6A. A subset of these modified peptides are quantified across hundreds of single cells, which enabled us to quantify differential abundance of phosphorylated peptides, phosphatases and kinases across cell types, Fig. 6B. Interestingly, the structural subunit A (PPP2R1A) and the regulatory subunit B (PPP2R3A) of the Serine/threonine-protein phosphatase 2A exhibit different abundance patterns across the cell types, suggesting different compositions of this protein complex across cell types. Many phosphorylation sites also exhibit differential abundance, including talin-1, which may affect the linkage of integrins to the actin cytoskeleton.

**Figure 6.**
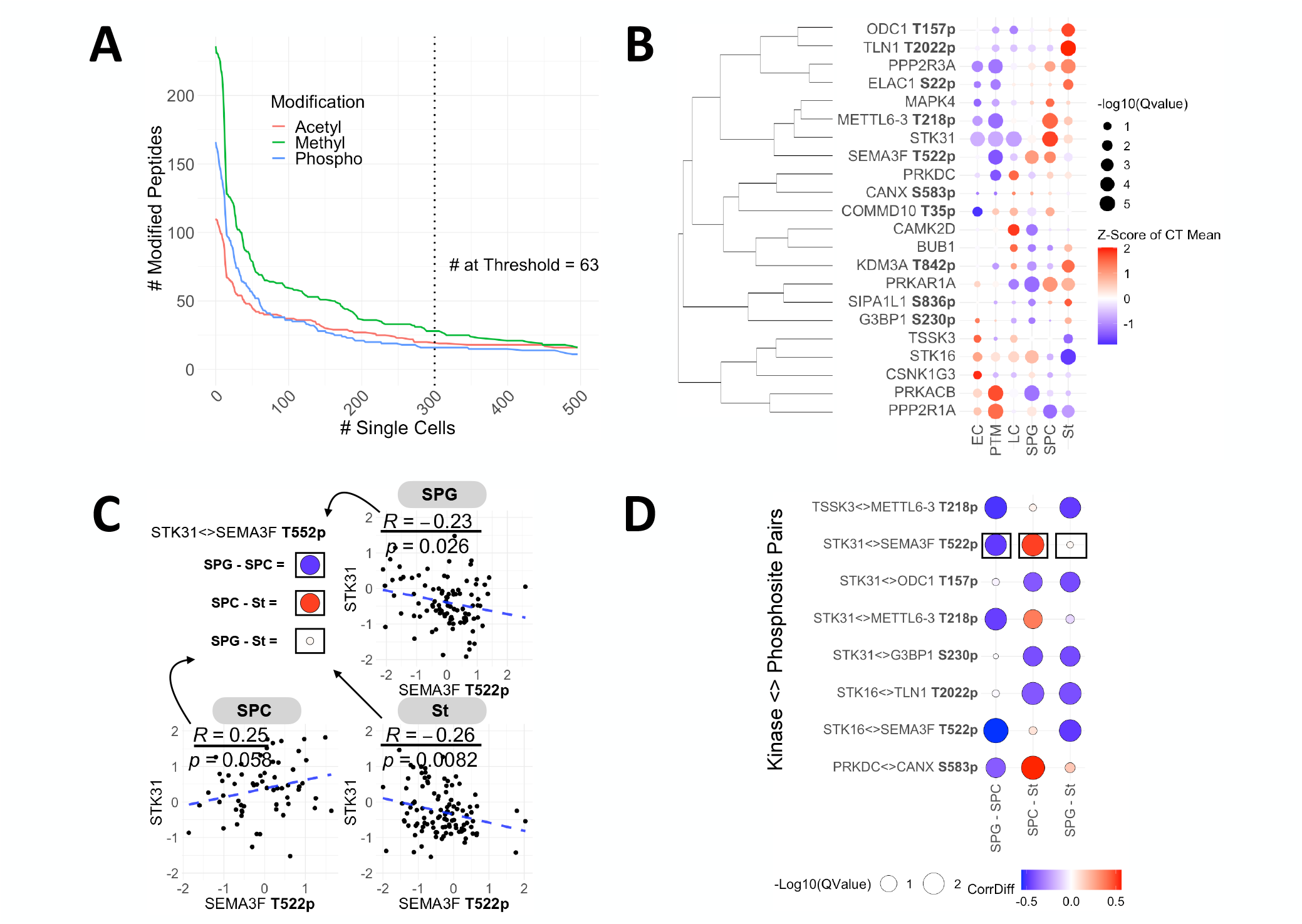
Covariation of post translationally modified peptides across single cells. **A** Cumulative distribution of PTMs across first 2 shotgun datasets, axis clipped at 500 cells. **B** Cluster level abundances of kinases and phosporylated peptides. **C** Statistically significant changes in correlations of phosphopeptides and kinases across the spermatogonial stem cell lineage, shown for 1 pair. **D** Shown for all pairs where differences in correlation are significant across the lineage.

To further examine phosphorylation signaling across cell types, we quantified the covariation of kinases and phosphorylated peptides in single cells, Fig. 6C,D. We observe that STK31 serine/threonine kinase 31 is significantly correlated to the abundance of phospho threonine 522 in SEMA3F, and the sign of this correlation changes across cell types, Fig. 6C. Similar changes in the sign of correlations are observed for multiple other pairs of kinases and phosphosites as shown in Fig. 6D. These include the phosphorylation of threonine 218 of METTL6, a tRNA methyltransferase, which can be a mechanism contributing to post-transcriptional regulation.

## Discussion

We introduce *BayesPG*, a Bayesian hierarchical model built to estimate reliability and quantify post transcriptional regulation. It leverages peptide and transcript levels measured across replicates and technology platforms to account for the technical variation and infer the biological variation. This is a major advance over previous analysis of RNA and protein levels from the same population of single cells^37^ that did did not model reliability. Modeling reliability enabled correcting for technical factors and estimating protein to mRNA relationships across cell types (Fig. 2D), highlighting the biological regulation. This allowed us to identify thousands of gene products and GO groups as candidates for significant post transcriptional regulation.

Our results indicate that only about half of the protein variance across cell types is captured by the corresponding RNA variation, Fig. 2. This fraction vary significantly across different functional groups of proteins, with some groups having low RNA-protein concordance and other much higher, Fig. 5. These estimates are in the same range as the estimates for variation across different human tissues^16,18^; They are lower than than the estimate for cells stimulated in vitro^**jovanovic2015dynamic**^, which likely reflects in part differences between steady-state and dynamic response to acute stimulation. Crucially, *BayesPG* allowed us to perform this resolution with cell-type resolution and discover regulation that is masked by tissue-level averages.

Our approach has some limitations. First, by modeling at the cluster-level, *BayesPG* does not capture important cell-to-cell variation in rPTR within clusters^38,39^. Second, the protein and RNA data came from different human subjects, which contributes variability to our estimates, which nonetheless is smaller than the differences between cell clusters, Extended Data Fig. 1. Third, gene-level scaling causes fold changes within each modality to shrink toward one another, which may yield more conservative posterior rPTR estimates. Fourth, cell type assignments may contain errors that lead to biased and/or noisier estimates of rPTR^40^. Such errors increase the uncertainty of the posterior distributions, and future developments may improve cell type assignment and error modeling. Finally, we assume that unobserved values are “missing at random”^41^. By modeling the nonignorable mechanisms that govern missingness^9^, we may ultimately infer more accurate estimates of rPTR.

While *BayesPG* focused on quantifying regulation across clusters, our data also allowed within cluster analysis of both proteins and post translational modifications. Specifically, we found significant covariation between kinases and phosphorylated sites that changes across cell types, Fig. 6. Analysis of protein covariation across single cells and nuclei has allowed inferring regulatory mechanisms of nuclear transport^42^, and it can similarly enable more regulatory inferences in complex tissues that the one we analyzed. The rigorous and systematic inference by *BayesPG* provides a stepping stone towards extending such regulatory inferences from single-cell proteogenomics data^19^.

## Methods

### Sample preparation

Testis tissues were dissociated as previously described in *Shami et al*^23^, single cell suspensions were preserved in a 90%FBS:10%DMSO solution. The samples were prepared for MS analysis using Nano-ProteOmic sample Preparation (nPOP) in 5 separate batches (3 x SCoPE2, 1 x pSCoPE, 1 x plexDIA) as described by Leduc *et al*.^27^. For each batch, an aliquot of sample was taken from stock tubes, washed in 10mL of PBS and spun down at 600g for 8 minutes. The cells were resuspended in PBS, filtered through a 40 *µM* cell strainer and spun down again. The cells were then resuspended in PBS to obtain a final concentration of 300 cells/*µL* and used for sample preparation. The first four sample preparations were done using an isobaric carrier channel in the SCoPE2 style of sample preparation using the TMTpro 18plex reagent (The first two channels were used for carrier, reference with the following 2 channels left empty, 14 channels were used to label single cells). The carrier and reference channels were prepared as a single batch as outlined in Petelski *et al*^43^. The carrier was benchmarked to be equivalent to 25ng of peptides. The final sample preparation for plexDIA analysis^25^ was also performed with nPOP, single cells were labelled with mTRAQ channels Δ0, Δ4 and Δ8. Prepared sets of single cells per sample prep batch were transferred to 384 well plates, dried down in a speedVac vacuum evaporator, sealed and stored at -80C. Prior to MS analysis, the SCoPE2 sets were resuspended in 1.05*µL* of 0.1% formic acid (buffer A), while the plexDIA sets were resuspended in 1.05*µL* of formic acid containing 0.015% DDM (N-Dodecyl B-D-maltoside).

**Table 1.**
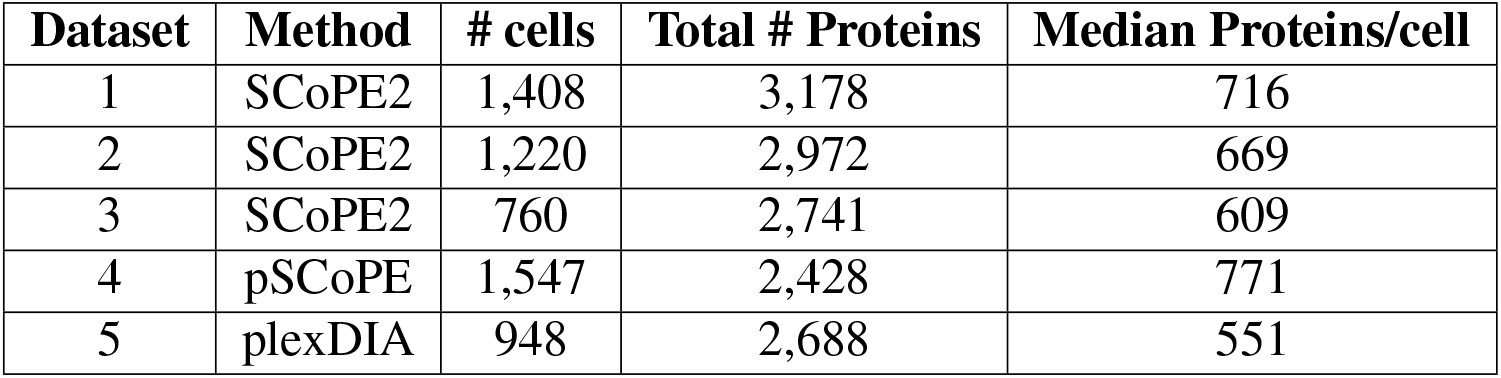
Overview of proteomic datasets. Summary data is provided using DART-ID upgraded outputs for the DDA data and match between runs enabled outputs for the DIA data. All data was filtered at 1% FDR.

| Dataset | Method | # cells | Total # Proteins | Median Proteins/cell |
| --- | --- | --- | --- | --- |
| 1 | SCoPE2 | 1,408 | 3,178 | 716 |
| 2 | SCoPE2 | 1,220 | 2,972 | 669 |
| 3 | SCoPE2 | 760 | 2,741 | 609 |
| 4 | pSCoPE | 1,547 | 2,428 | 771 |
| 5 | plexDIA | 948 | 2,688 | 551 |

### Data acquisition

#### Peptide separation

For all sample preparations, 1*µL* of sample was loaded onto 25cm x 75*µM* IonOpticks, C18 columns. We used Odyssey series with a liquid-liquid junction for the Ultimate 3000 and Aurora series with captive spray insert and nanoviper junctions for the the Vanquish Neo. The peptide separations for all samples was performed at flow rate of 200nL/min.

The sets run on the Ultimate 3000 consisted of a 100 minute total run time per sample. Samples were loaded onto the separation column for 20 minutes, followed by a linear gradient from 8%B buffer B (80% acetonitrile and 0.1% formic acid) to 26% buffer B for 63 minutes. The column was then was at 95%B (5 minutes total) and equilibrated at 4%B for 20 minutes.

The sets run on the Vanquish Neo used about 30 minute total run time per sample, which was loaded in 1.2*µLs* via direct injection. Samples were loaded onto the separation column by maximizing flow rate to maintain LC system back pressure at 1450 bar (with a max ramp of 900 bar/second). The separation was performed using a dynamic active gradient (15 minute total) which was as follows: 2.5%B to 6.5%B over 0.2 minutes, to 11.5%B over 0.9 minutes, to 21.0%B over 3.1 minutes, to 31.5%B over 6.2 minutes, to 40%B over 2.8 minutes, to 55%B over 1.7 minutes. The column was washed at 95%B for 4.65 minutes. Equilibration was performed similar to the loading for a volume that is 4 times the void volume of the column.

#### Acquisition of mass spectra

For the SCoPE2 sets all spectra were acquired in shotgun DDA mode using a Thermo Scientific Q-Exactive mass spectrometer from minutes 20 to 95 of the LC method. The electrospray voltage of 1700 V was applied at the liquid–liquid junction of the analytical column and transfer line. The temperature of the ion transfer tube was 250 °C, and the S-lens RF level was set to 80. After a precursor scan from 450 to 1600m/z at 70,000 resolving power, the top 4 most intense precursor ions with charges 2 to 4 and above the AGC min threshold of 20,000 were isolated for MS2 analysis via a 0.7 Th isolation window with a 0.3 Th offset. These ions were accumulated for at most 500 ms before being fragmented via HCD at a normalized collision energy of 33 eV (normalized to m/z 500, z = 1). The fragments were analyzed at 140,000 resolving power. Dynamic exclusion was used with a duration of 30 s and a mass tolerance of 10 ppm.

The prioritized data acquisition for pSCoPE used the same instrument parameters as those described for shotgun acquisition, but the precursors were selected for isolation and fragmentation using MaxQuant.Live and inclusion lists described below.

The plexDIA sets were acquired in DIA-PASEF mode using a Bruker TIMS-TOF SCP. The duty cycle was optimized for frequent precursor sampling. Specifically, it consisted of 8 PASEF frames with 26 Th MS2 windows (1 Th overlaps). An MS1 scan was taken every 2 PASEF frames, resulting in 4 MS1 scans per duty cycle. The MS1 scan range was 100-1700 m/z, while MS2 scan range was 300-1000 m/z. The 1/K0 range was between 0.64 and 1.20 and collision energy was set at 20eV at 1/K0 = 0.60 and 59eV at 1/K0 = 1.60, collision RF was set to 2000 Vpp. The ramp and accumulation times were 100 milliseconds and estimated duty cycle time is 1.28 seconds.

### Searching acquired mass spectra

The SCoPE2 style data was searched using MaxQuant against a protein sequence database that included all entries from the human Uniprot database (UP000005640; 101,014 entries) and known contaminants. The MaxQuant search was performed using the standard workflow, which includes trypsin digestion and allows for up to two missed cleavages for peptides with 7 to 25 amino acids. Tandem mass tags (TMTPro 18plex) were specified as fixed modifications, while methionine oxidation and protein N-terminal acetylation were set as variable modifications. For identifying post translational modifications (PTMs) Acetyl (K), Phospho (STY), Methyl (KR) were set as variable modifications instead, these searches were limited to only the sets acquired using shotgun DDA (first three sample preparations). Carbamidomethylation was disabled as a fixed modification, because cysteines were not alkylated. Second peptide identification was also disabled. The calculation of the peak properties was enabled. All peptide-spectrum matches (PSMs) and peptides found by MaxQuant were exported to the evidence .txt files. The confidence in the PSMs was further updated using DART-ID, which is a Bayesian framework for increasing the confidence of PSMs that were consistently identified at the same retention time with high-confidence PSMs for the same amino acid sequences^44^. For the standard search, DART-ID was carried out within each sample preparation batch independently. For the PTM search, we carried out DART-ID separately for each modification, across all 3 sample preparation batches.

The updated data were filtered at 1% FDR for both peptides and proteins as described by Petelski *et al*^43^.

The plexDIA data was searched using DIA-NN (version 1.8.1)^45^. A predicted spectral library was generated by in silico labeling a SwissProt (2021-10-05 release ; 20,347 entries) human database with mTRAQ on each trypsin-digested peptide. The ion mobilities (IMs) for each mTRAQ-labeled precursor predicted by DIA-NN were refined using mTRAQ-specific predictions. The refined predictions were made using an ensemble of five deep neural networks, the loss function was the mean absolute error for predictions weighed by the MS1 profile correlation. The library was then used to search a small sample (2.5ng peptides, split across all three mTRAQ labels) and the search output was used to generate an empirical library (5,300 protein groups, 52,000 precursors). The single cell sets were searched with DIA-NN using this empirical library. Peak height was used for quantification with a scan window of 5, mass accuracy of 15 ppm and MS1 accuracy of 10 ppm. Peak translation and MBR were enabled, and search outputs were filtered at 1% Q value.

### Inclusion list generation for pSCoPE

To generate the inclusion list for pSCoPE analysis, we first analyzed a 5x carrier sample (∼125ng of peptides) using DIA Methods 1 and 2 which are outlined in Supplemental Table S3 of Huffman et al. The DIA data acquired for the TMT labeled carrier samples were only used for generating accurate retention times for precursors and not for any quantification. An empirical spectral library was generated by searching DIA spectra together with shotgun DDA spectra from sample preparation batches 1 and 2 using FragPipe^46,47^ (FragPipe version 17, MSFragger version 3.4). The library was generated using the DIA Speclib quant workflow and TMT was added as a fixed modification. The spectral library was subsequently used to search a 1x carrier sample acquired with DIA method 1 to obtain accurate retention times for precursors. The 11,931 peptides obtained from this search were split across four priority tiers (tier 0 = used for retention time alignment only, 4 = highest priority tier). Tier 4 consisted of peptides that had previously been identified as unique markers for each our 6 celltypes. The remaining peptides were assigned to tiers such that higher intensity, more confidently identified peptides were put in higher tiers, we did not target peptides mapping to proteoforms. We tested this list on a 1x carrier injection and searched the acquired DDA spectra using MaxQuant (against the SwissProt sequence database; 2021-10-05 release ; 20,347 entries). We readjusted and iterated on our inclusion list to optimize for precursor identifiability as outlined in Huffman et al. Fill times were increased for a subset of the marker peptides in the top tier which had a lower identification rate.

### Data Processing

For the SCoPE2 data, MaxQuant output for each sample preparation batch was independently processed (using the SCoPE2 pipeline) to yield their associated protein x cell matrices. The only deviation was that the sample loading normalization was performed using the intersected median across all cells as opposed to that of each cell. Specifically, the median value for each feature across all cells was assigned to a reference vector. The median difference between the non missing values from each cell and the reference vector were then used to scale all feature values for that cell. Protein values were obtained by selecting the median across peptides which map to it, including those which map to proteoforms. Batch correction (using ComBat, at the protein level) within a sample preparation batch was only performed for a subset of runs in the first batch which had systematically higher intensities than all other runs.

The plexDIA data were processed using the QuantQC package^48^. The MS1 area of peptides in each cell was normalized using the intersected median normalization as above. Cells were filtered for quality by comparing the median coefficient of variation for proteins with more than 3 peptides within each cell to that of negative controls. Protein level quantification was estimated from the median peptide relative levels, as done in the SCoPE2 pipeline. Peptides that mapped to multiple proteins were removed from the analysis. Label biases for mTRAQ were corrected using ComBat^49^.

### Cross modality dataset alignment

#### Pre-processing and feature selection

The protein x cell matrices from each batch were subset to the intersect of proteins across batches and merged. The merged matrix was then batch corrected using ComBat^49^. The unimputed, batch corrected matrix was used to selected the gene product space within which cross modality alignment was carried out.

The single cell mRNA-Seq datasets were processed using Seurat^50^, version 4.1.1. The data from Sohni *et al*^22^ consisted of a count matrix corresponding to either donor which was then filtered for technical artifacts using mitochondrial reads, number of features. *Shami et al*^23^ had the data only available as one count matrix which had already been filtered for artifacts. Sctransform^51^ was used to normalize each matrix independently using 5000 variable features (all features were output post normalization) and regressing out the contribution of mitochondrial read percentage on the variance. Each matrix was then subset to the intersect of all gene products across protein and mRNA matrices respectively. The subset matrices were then integrated using the integration anchor based workflow implement in Seurat v3, a detailed implementation can be found in Stuart *et al*, 2019^52^.

To select the feature space for dataset alignment we first computed gene product correlation matrices within each modality. We then correlated each corresponding vectors of pairwise correlations for RNA and proteins. We selected the subset of gene products with correlations above the median correlation, and recomputed the correlations between correlation vectors. The RNA and proteins with correlations above the median were selected to carry out dataset alignment.

#### Alignment and validation

Dataset alignment was carried out using integrative non-negative matrix factorization (iNMF) as implemented in the LIGER package, Rliger^28^. As we require non-negative matrices, the mean centered and log2 transformed protein x cell matrix was exponentiated. For the purposes of the alignment only, we use the imputed and batch corrected (for sample preparation batch). The mRNA data were used without any processing. The data were subset to the previously selected alignment space and mRNA data were size normalized and scaled without centering. The iNMF was performed using 10 meta-gene factors (k) and regularization parameter (*λ*) of 5. These parameters were selected by optimizing for dataset alignment and agreement scores, as calculated in LIGER. The shared factor neighbourhood graph was then jointly clustered, quantile normalized and louvain clustering was carried out to further refine and obtain final cross modality clusters. Cells within each louvain cluster were assigned to the cell type with the highest proportion of annotated mRNA single cells. We focused our analysis on 6 celltypes: Endothelial cells, Peritubular Myoid Cells, Leydig Cells, Spermatogonia, Spermatocytes, Spermatids. Since the mRNA datasets had annotations at different levels (cell type vs sub celltype), all annotations were collapsed to the celltype level; for example, round and elongating spermatids were considered spermatids.

We first evaluate our alignment procedure by assessing the agreement of gene products within the alignment feature space. For each cell type, gene product fold changes are averaged within each modality and then standardized using Z-scores. We then compute the cross-modality correlation for each cell type. To visualize clustering of celltypes transferred to the proteomic data we carried out principal component analysis (via Eigendecomposition) on the unimputed, batch corrected matrices in the alignment feature space. The first two eigenvectors were plotted and individual cells were colored by their celltypes. To quantitatively assess the clustering, we computed the Mahalanobis distance in the space defined by Eigenvectors 1 and 2. For each Louvain cluster, we calculate a distance ratio. The ratio compares the median distance of cells within a cluster to the median of their distances to all cells in a different cluster, pairwise, for all clusters.

### Analysis of post translational modifications

To analyze PTMs across single cells, we start our analysis by refining our list of PTMs (which were filtered)at 1% global FDR). We compute a local, modification specific FDR using the DART PEP and control the rate of false discoveries at 5%. We limited our analysis to post translationally modified peptides that are observed in at least 300 cells across the datasets. For our analysis of associations between phosphorylated peptides (phosphopeptides) and kinases, we further subset to kinases which had at least 1000 pairwise observations across all phosphorylated peptides and vice versa. This yielded a set of 13 kinases and 11 phosphopeptides.

To evaluate associations across all celltypes we first collapsed the features’ relative levels across cells to the mean per cell type. We then compare the mean for each feature per celltype to the other celltypes and compute p values using the Wilcoxon rank sum test and correct p values for the multiple hypotheses tested using the Benjamini Hochberg procedure. To visualize the associations we standardized celltype abundances using Z-scores and hierarchically clustered the features (euclidean distances and complete linkage) to produce a dendrogram.

To test the significance of the difference between correlation across celltypes we employed Fishers Z transformation to allow us to compare pairwise correlations. Specifically, for each correlation pair the Fishers Z transform was applied to the correlations and the difference was computed between these values. The standard error of the differences was computed as follows: 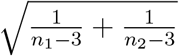 where the sample size ’n’ was set to 100 for each condition (n1, n2, n3). A Z score was then calculated by dividing the absolute difference by the standard error and a two-tailed p-value was obtained using the normal distribution. The p values were then corrected for the multiple hypotheses tested using the Benjamini Hochberg procedure.

### Modeling relative protein to mRNA ratios with *BayesPG*

We model for transcript count totals and peptide-level intensities across cell-type clusters using our novel method, *BayesPG*. Let 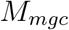 represent the sum of transcript counts across cells assigned to cell type *c* for gene *g* and mRNA data set *m*. Let 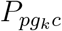 represent the average log_2_ peptide intensities for protein data set *p* across cells assigned to cell type *c* for peptide *k* associated with gene *g*.

We assume that cells are correctly associated with their corresponding cell type, which depends on the performance of data set alignment and cell type identification, and that peptides are properly identified during the data collection process. For each data set, we assume that gene product abundances are conditionally independent given model parameters. Another critical assumption for this model is that *M*_*mgc*_ and 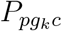 are independent replicates corresponding to a shared underlying biological process. In particular, transcript counts for gene *g* among cells assigned to cell type *c* have some shared log_2_ transcript abundance, *µ*_*gc*_, and peptide-level averages have shared underlying log_2_ protein intensity *µ*_*gc*_ + *r*_*gc*_, where *r*_*gc*_ is defined as the log_2_ relative ratio of protein to mRNA. The log_2_ relative protein-to-mRNA ratio, *r*_*gc*_, reflects the degree of post transcriptional regulation affecting gene *g* within cell type *c* relative to the average transcript or peptide intensity corresponding to gene *g*. If there is no post transcriptional regulation, the amount of protein synthesized will be directly proportional to the amount of mRNA transcribed and *E*[*r*_*gc*_] = 0. While empirically computed rPTR is subject to high measurement noise and missingness, posterior rPTR samples estimated with *BayesPG* are more robust. This is because of the use of replicates across peptides associated with the same protein and data sets, which offers the opportunity to estimate the effect of distinct sources of variability– that which is associated with biological processes, technical noise across peptides associated with the same protein, and technical noise across data sets within the same modality.

### Cluster level observations

Figure 1 displays summary information for cluster-level transcript and peptide data. Protein consistency is computed using the correlation of across-peptide *z*-transformed log_2_ cluster-level averages for two sets corresponding to each protein with 2 or more peptides observed, where peptides are randomly assigned to each set for each of such proteins. To compute data set agreement, z-transformed log_2_ cluster-level averages are computed for each protein, cluster, and data set and are correlated across clusters between each pair of data sets.

### Likelihood

#### mRNA

We model the sum of transcript counts across cells assigned to each cell type for each individual gene, *M*_*mgc*_ using a Negative Binomial with mean with mean 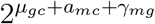 and dispersion *ϕ*_*g*_:

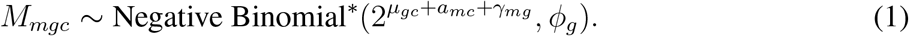

To ensure that log_2_ transcription rate, *µ*_*gc*_, is defined relative to its gene-level average, we use parameter *γ*_*mg*_ to represent the average transcript level in gene *g* across all cell types in each mRNA data set *m*, for data set associated biases that may affect transcript counts independently for different genes. Similarly, *a*_*mc*_ cell-type normalization term, that accounts for data set associated systematic effects biasing cell types. The overdispersion term *ϕ*_*g*_, accounts for the high variability associated with modeling transcript count data, as a result of non-biological sources of variability and biological-source related variability, including transcriptional bursts, where transcription occurs in “pulses”^53^.

#### Protein

We model 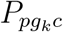, the average log_2_ peptide-level intensity across cells associated with each cell type for proteins corresponding to each gene *g* conditional on corresponding mRNA levels. Specifically, we assume

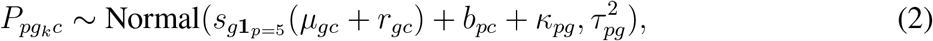

which assume each peptide is an independent replicate from the protein to which it forms.

We model gene-level average protein intensities with *κ*_*pg*_ in order to consider biological process parameters *µ* and *r* relative to the average protein-level intensity associated with each gene, and account for gene-level systematic effects associated with each data set. Similarly *b*_*pc*_ represents the cell type, data set normalization term.

We use scale parameter 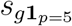 to account for ratio compression in individual proteins and data collection protocols. This ensures that biological process parameters are on the same scale across modalities. This scaling parameter is shared across (p)SCoPE-based data sets, *p* = {1, …, 4}, and modeled separately for the plexDIA-based data set, 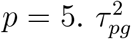 represents sampling variability for peptide-level averages associated with protein *g* and data set *p*.

#### Prior distributions

Biological process parameters *µ*_*gc*_ and *r*_*gc*_ are assigned mean-zero normal priors, since both *µ*_*gc*_ and *r*_*gc*_ are modeled respective to gene-level means. A priori, this choice shrinks estimates toward a null of no fold-change differences across cell types and no post-transcriptional regulation.

The scale parameters, 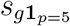 are assigned inverse Gamma(2, 1) distributions. This prior is also shared across measurement techniques, (p)SCoPE and plexDIA, so that no difference in scaling is assumed a priori.

We introduce gene product normalization terms *γ*_*mg*_ and *κ*_*pg*_ for mRNA and protein data sets, respectively. Similarly *a*_*mc*_ and *b*_*pc*_ represent cluster-level normalization parameters. *γ*_*mg*_ and *a*_*mc*_ are given Normal(0, 10) priors and *κ*_*pg*_ and *a*_*pc*_ are assigned Normal(0, 1) priors.

We model protein sampling variability 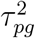 using a hierarchical structure, where 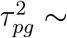 Normal 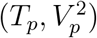 and Normal^+^(0, 1) priors are set on *T*_*p*_ and *V*_*p*_. This allows for sampling variability to be modeled independently within each protein and data set, while allowing for similarity across the variances of proteins observed in the same data set.

We model mRNA using the Negative Binomial distribution with alternative parameterization. We use *ϕ*_*g*_ to govern overdispersion for each gene and 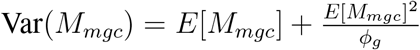 We place a Normal^+^(0, 1) prior on 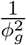 which puts more mass a priori on fits with no overdispersion.

Table 2 displays a full specification of parameters present in the model as well as corresponding prior distributions. All parameters parameters are assumed to be independent a priori.

**Table 2.** Model parameters and corresponding prior distributions.

| Parameter | Interpretation | Prior Distribution |
| --- | --- | --- |
| $\mu_{gc}$ | Underlying $\log_2$ transcription rate | Normal(0, 1) |
| $r_{gc}$ | Log relative protein-to-mRNA ratio (rPTR) | Normal(0, 1) |
| $a_{mc}$ | mRNA cluster-level normalization | Normal(0, 10) |
| $b_{pc}$ | Protein cluster-level normalization | Normal(0, 1) |
| $s_g \mathbf{1}_{p=5}$ | Protein scaling parameter | $1/s_g \sim \text{Gamma}(2, 1)$ |
| $\gamma_{mg}$ | mRNA gene-level normalization | Normal(0, 10) |
| $\kappa_{pg}$ | Protein gene-level normalization | Normal(0, 1) |
| $\tau_{pg}^2$ | Protein sampling variability | $\tau_p^2 \sim \text{Normal}(T_p, V_p^2)$ |
| $\phi_g$ | mRNA overdispersion | $1/\phi_g^2 \sim \text{Normal}^+(0, 1)$ |

### Inference and model checking

We fit *BayesPG* using the probabilistic programming language Stan^30^, which uses No-U-Turn Sampling (NUTS) to perform fully Bayesian inference. We generate 8,000 posterior samples across 10 chains and verify that the chains have sufficiently mixed by examining Rhat diagnostics.

To evaluate model fit we use posterior predictive model checking. Specifically, we compare statistics computed from observed data to those computed from simulated data generated from the posterior predictive distribution. In Figure 5A, we examine the posterior predictive coverage for 1) across cluster fold changes of mRNA and protein and 2) the across cluster variance of mRNA and protein. In Figure 5B we show that the distribution of across cell type correlations in the generated data is broadly consistent with the distribution for the observed data. We note that the observed across cell type mRNA, protein correlations is simply the across-modality data set agreement explored in Figure 1A.

### Significance testing

#### Selecting Gene Products with Significant rPTR

To identify genes with statistically significant rPTR, we compare the rPTR in each gene and cell type to zero. We compute 95% posterior intervals across *i* for each gene and cell type. A gene-cell type pair is reported “significant” if the 95% posterior credible interval excludes 0.

For gene-level testing, we define the posterior exclusion probability (PEP) as,

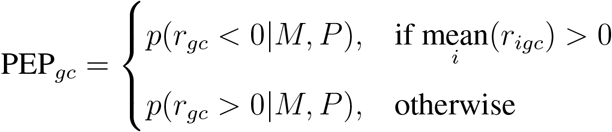

where we define *p*(*r*_*gc*_ *<* 0|*M, P*) as the posterior probability *r*_*gc*_ *<* 0 given mRNA and protein measurements. *p*(*r*_*gc*_ *<* 0|*M, P*) is approximating using Monte Carlo samples. We compute the expected proportion of false discoveries, FDR_*gc*_, as the cumulative mean of the ordered PEPs across genes for each cell type.

#### Selecting significant gene ontology groups and protein complexes

To test rPTR in GO groups and protein complexes (collectively referred to as “groups”), we compare group averages of rPTR to the overall average rPTR across all cell types. In doing so, we identify groups for which rPTR in a given cell type is distinguishable from others. We first center *r* at the group-level using the average across clusters.

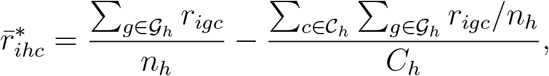

where *n*_*h*_ is the number of gene products associated with group *h, C*_*h*_ is the number of cell types observed for group *h*. If the 95% posterior intervals for 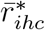 excludes 0, we declare that GO group significant for that cell type. We compute FDRs in the same way as gene-level testing using 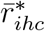 instead of *r*_*gc*_. We restrict testing to groups with at least 25% gene products observed so as to avoid test results that aren’t representative of a given group.

#### Selecting Significant Correlations in Gene Ontology Groups and Protein Complexes

We test GO groups and protein complexes for those with significant high and low across cluster mRNA-protein correlation. The same 25% observation threshold is placed on groups as described in the previous section. For each complex, we compute the correlation of *µ*_*igc*_ and *µ*_*igc*_ + *r*_*igc*_ across cell types *c* and retain the median of these correlations across gene products *g*. Let *ρ*_*ih*_ represent this correlation for each set of posterior samples *i* and group (GO or protein complex) *h*.

Next we define 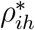, which allows us to test the complex-level correlations relative to the average,

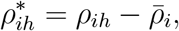

where 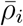 is the average of *ρ*_*ij*_ across groups *h*. We define 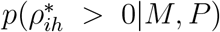 as the posterior probability that 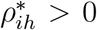. We approximate the posterior exclusion probability using Monte Carlo samples for each group,

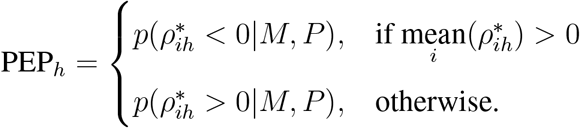

Next we compute the cumulative mean of PEP_*h*_ across groups *h* to identify significant complexes at a 5% threshold.

For each GO group and protein complex, we compute the median pairwise mRNA-mRNA and protein-protein across clusters correlation. For all unique pairs of genes *a* and *b* and each posterior draw *i*, we compute the correlation of *µ*_*iac*_ and *µ*_*ibc*_ across clusters *c*. Next, the median is computed for each set of posterior samples *i* across pairs of gene products associated with each group. Posterior summaries are computed across *i*. Similarly, for consensus protein draws, *µ*_*iac*_ + *r*_*iac*_ and *µ*_*ibc*_ + *r*_*ibc*_, we compute across clusters correlations and medians across pairs of gene products.

## Availability

Further documentation on the use of *BayesPG* as well as links to data repositories is available at scp.slavovlab.net. The code is is open source and freely available at https://github.com/SlavovLab/BayesPG.

## Acknowledgments

We thank Prof. Sue Hammoud for providing testis samples and helpful discussions. The work was funded by an Allen Distinguished Investigator award through The Paul G. Allen Frontiers Group to N.S., a Seed Networks Award from CZI CZF2019-002424 to N.S., an NIGMS award R01GM144967 to A.F and N.S., and a Bits to Bytes award from MLSC to N.S.

## Extended Data Figures

**Extended Data Fig. 1.**
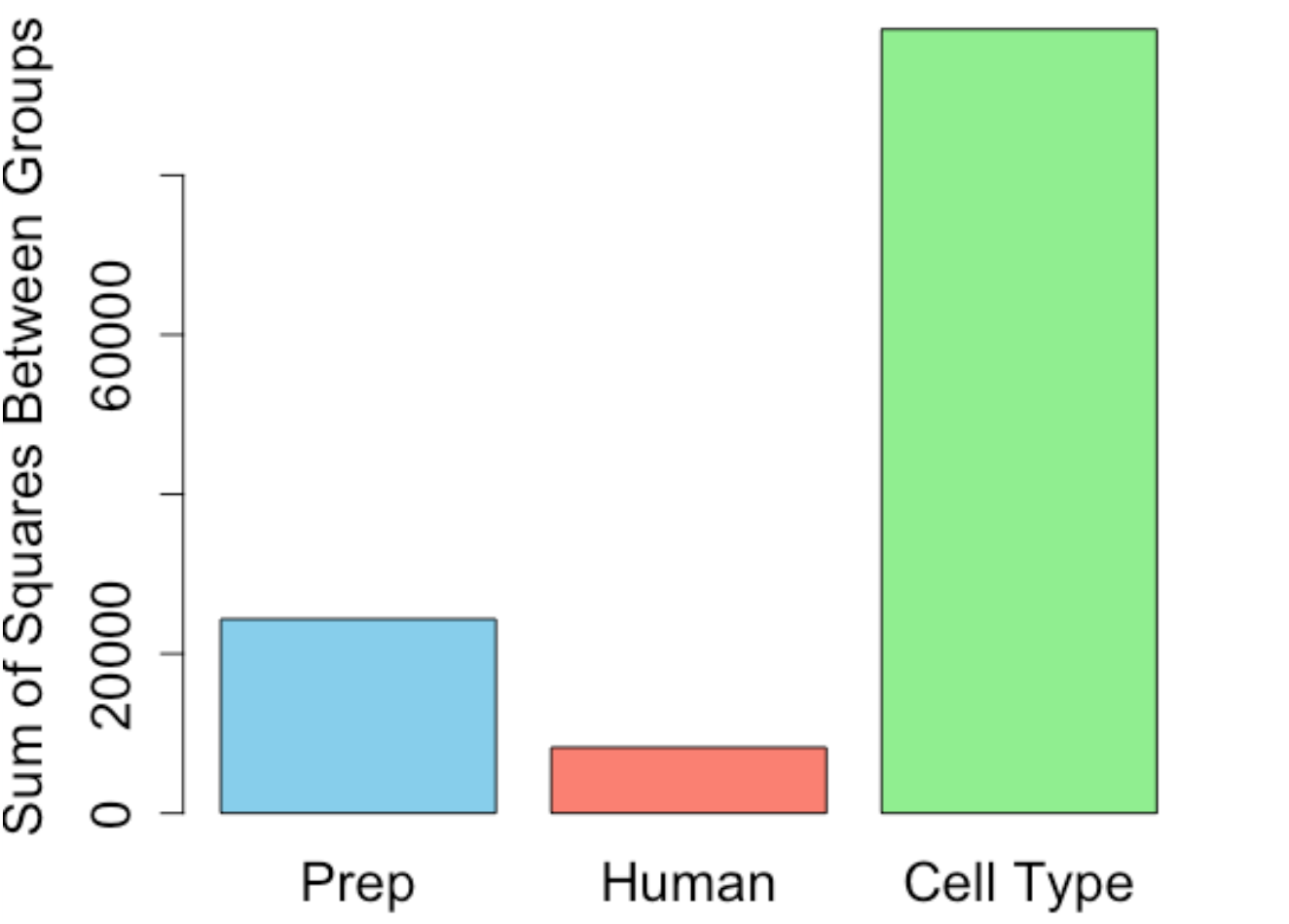
Comparison of the different sources of variation across mRNA-Seq datasets. The bar plot displays the sum of squares between groups for each covariate in the ANOVA model used for this comparison. Covariates included the prep (study from which data was taken), donor, and celltype (using original annotations). The analysis highlights the predominant influence of celltype on variance.

**Extended Data Fig. 2.**
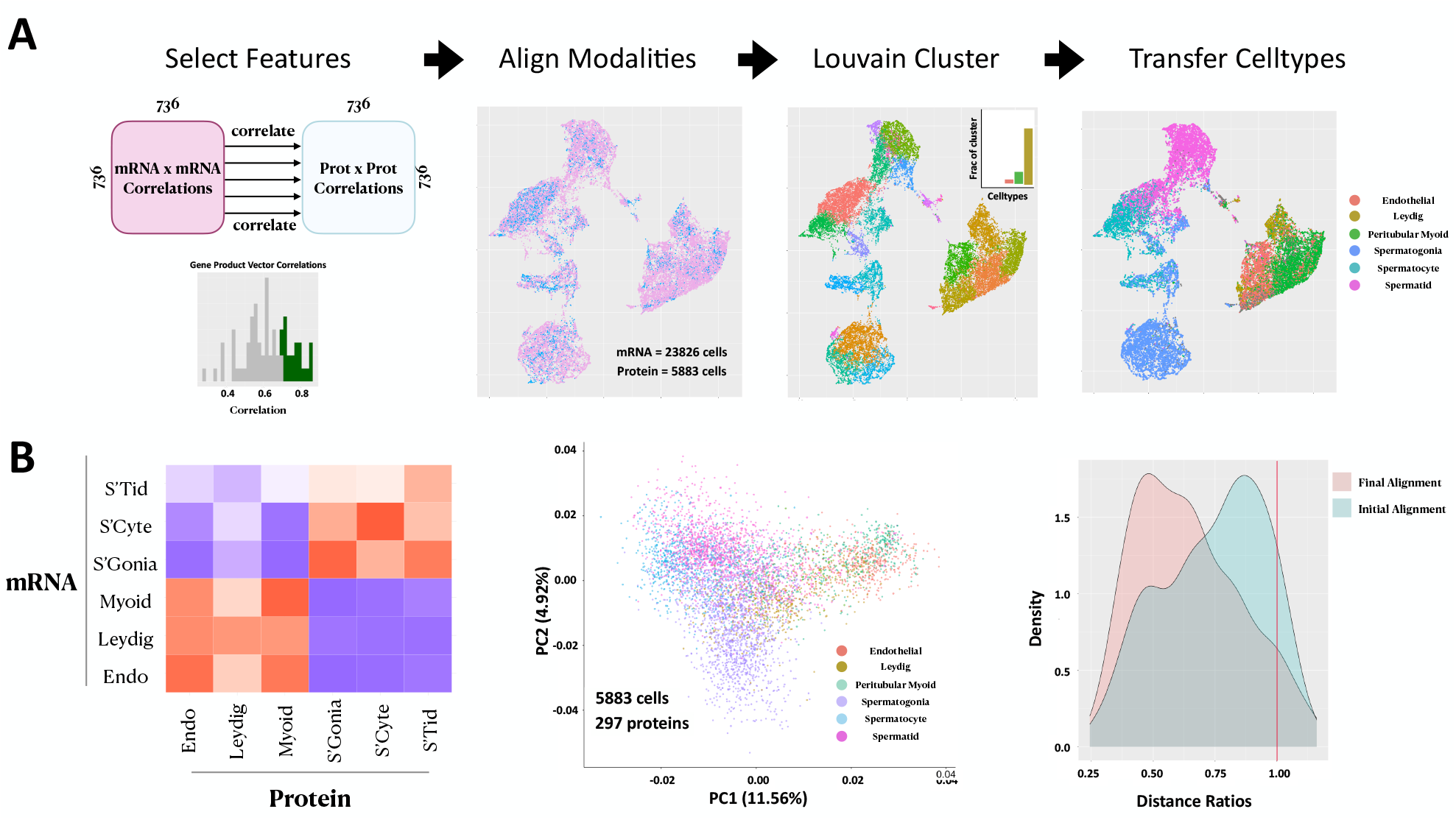
Data set alignment procedure. **A)** Alignment feature space is selected by correlating vectors of within modality correlations, highly correlated gene products are selected. Alignment is carried out using LIGER, the shared factor neighbourhood graph is subsequently Louvain Clustered and finally, each joint cluster is assigned to the cell type (using annotated mRNA data) with the highest fraction of cells within the cluster. **B)** Examples of metrics used to characterize alignment success. **Left)** Heatmap showing gene product level correlations aross modalities in the alignment feature space. **Middle)** Protein only cells visualized on Principal Components 1 and 2, colored by cell types assigned via alignment. **Right)** Distance ratios of clusters: the median Mahalanobis distance within a cluster divided by the median distance to each other cluster

**Extended Data Fig. 3.**
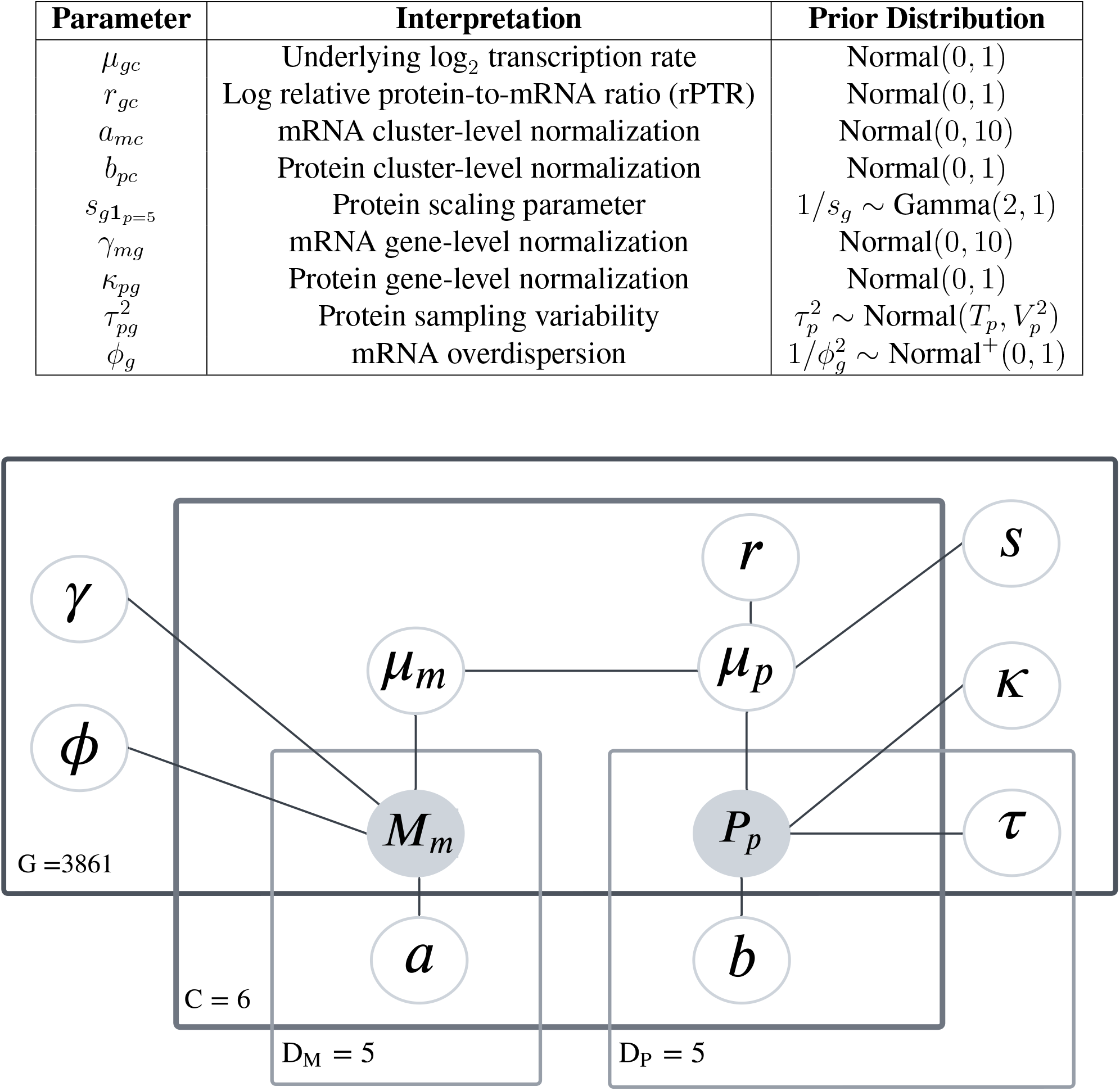
Plate diagram depicting model structure. *M*_*m*_ and *P*_*p*_ cluster-level observed values for the transcript and peptide data sets, respectively. *G* = 3861 refers to the total number of gene product pairs included in the model. Parameters within the corresponding plate are vectors indexed by gene product. *C* = 6 represents the 6 cell types for which data is modeled, with *µ*_*m*_, *µ*_*p*_ and *r* varying with gene product and cell type. *D*_*M*_ and *D*_*P*_ refer to the five datasets for each modality. Parameters within the *D*_*M*_ and *D*_*P*_ vary based on the plates within which they are nested.

**Extended Data Fig. 4.**
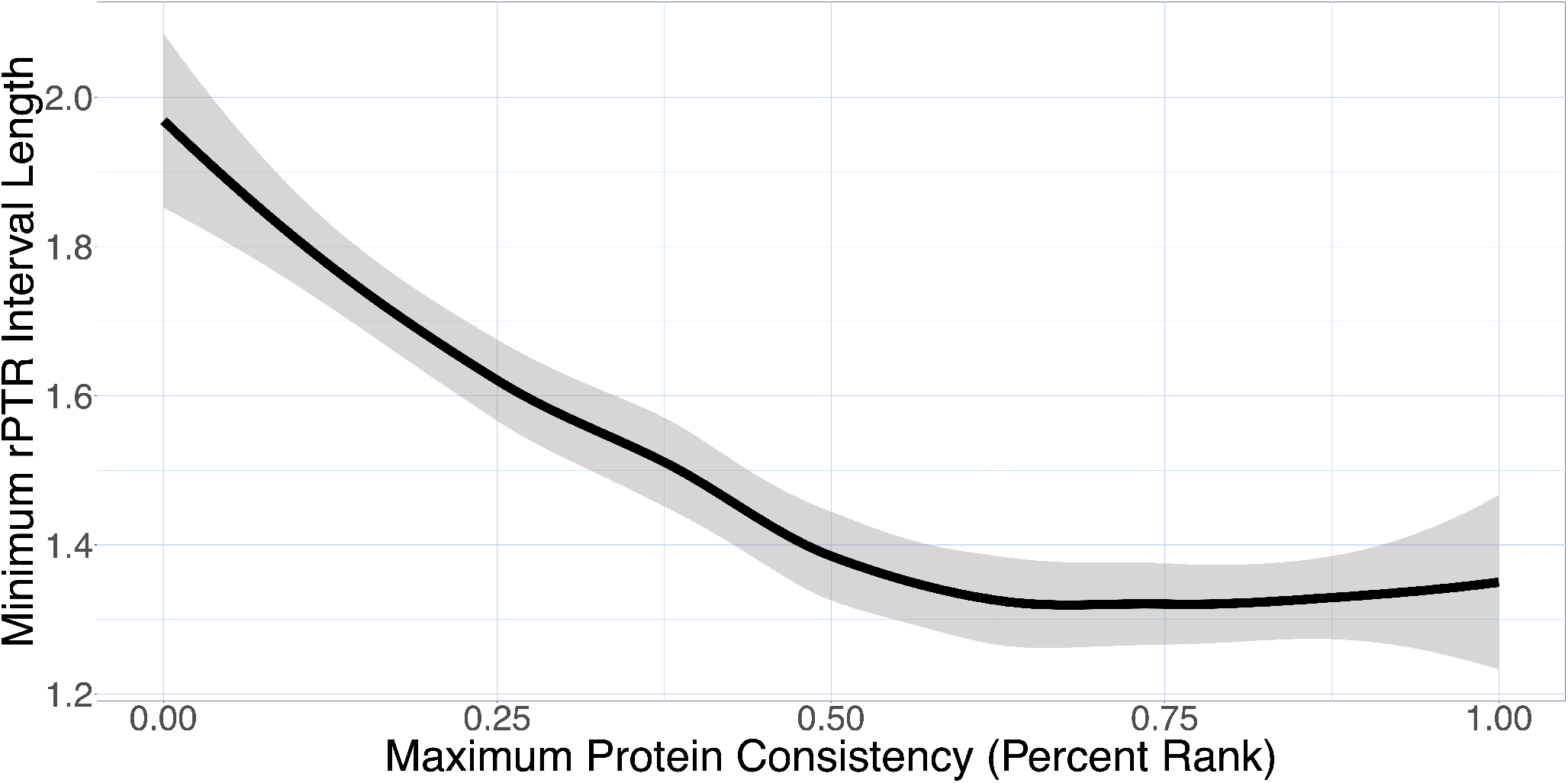
Minimum rPTR posterior interval width (y) by percent rank of maximum protein consistency (x). We compute interval lengths using 95% posterior intervals for rPTR for each gene product and cluster and use the minimum across clusters. Gene products with large disagreement between reliability metrics are excluded.

**Extended Data Fig. 5.**
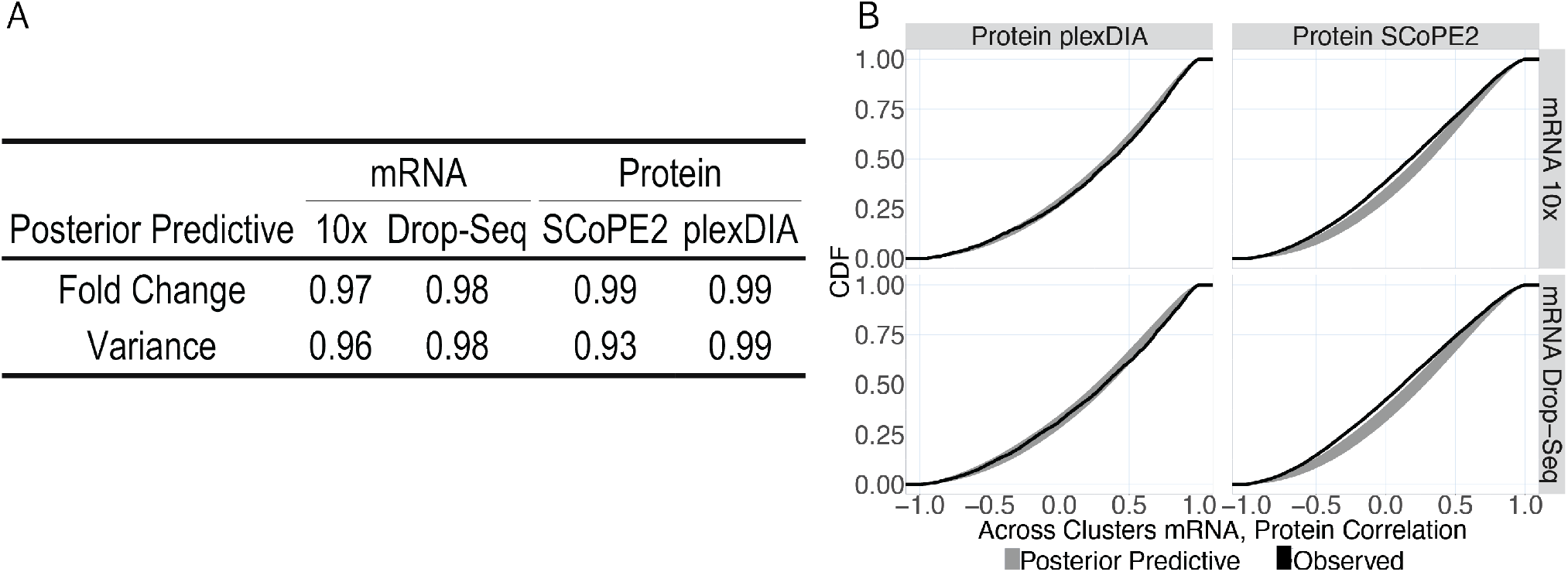
**A** Coverage and correlation of posterior predictive fold change and across clusters variance on observed values for each modality and data set. **B** Cumulative distribution function of posterior predictive across clusters mRNA-protein correlation, displayed in gray, and observed across clusters mRNA, protein correlations in black.

**Extended Data Fig. 6.**
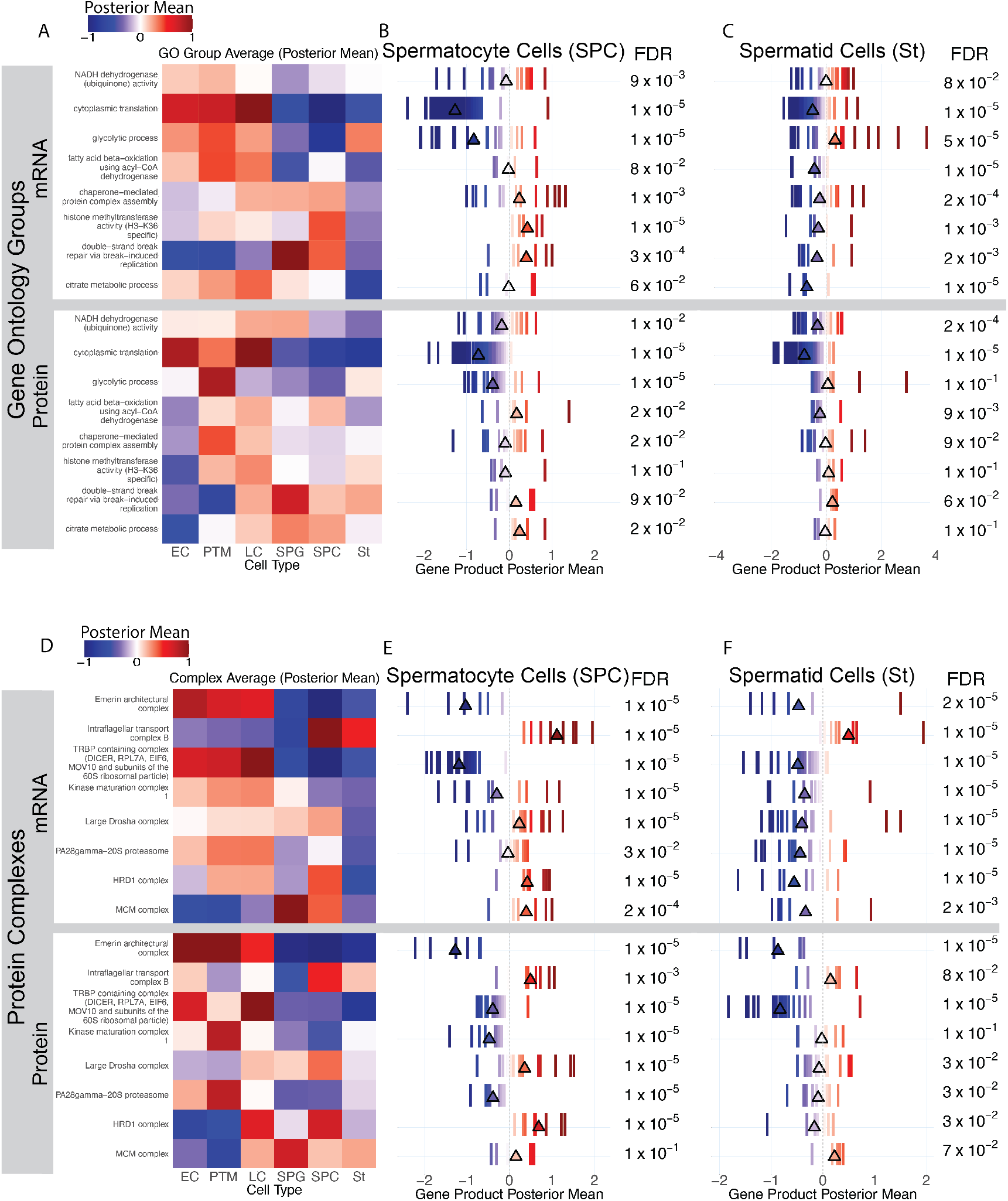
mRNA and protein levels for GO groups and protein complexes with significant rPTR from Fig. 3. The gene products and display is identical as in Fig. 3. **A-C** Results GO Groups, with RNA abundance (top) and protein abundance (bottom). **D-F** Results for protein complexes.

**Extended Data Fig. 7.**
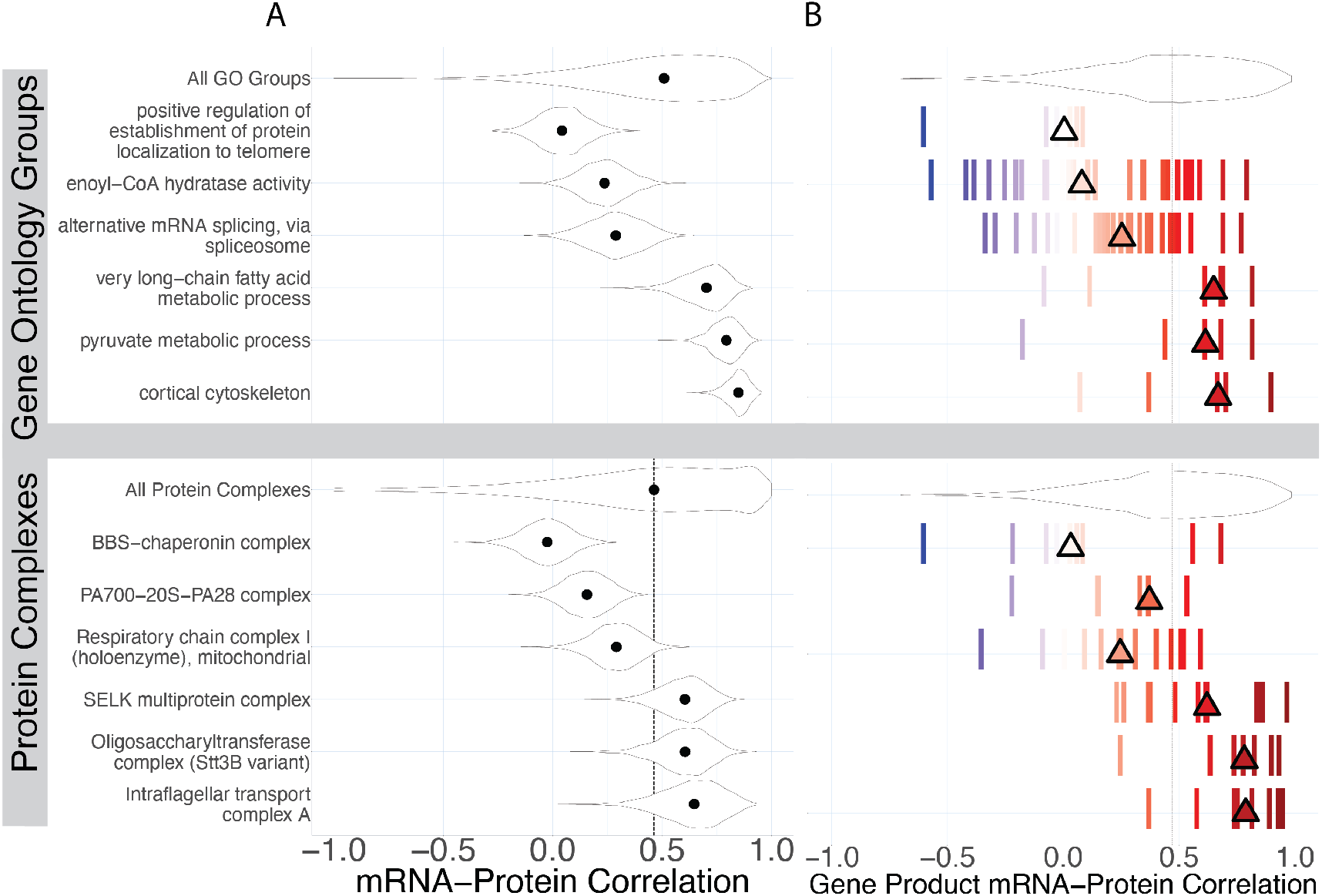
Additional posterior summaries for correlation test and groups highlights in Fig. 5. **A** Estimated posterior distribution of median mRNA-Protein correlation across gene products associated with each group. Points represent posterior median across posterior draws. **B** Posterior mean of mRNA-protein correlation for gene products associated with groups shown in **A**. Color also represents posterior mean, and triangles display median across gene products.

**Extended Data Fig. 8.**
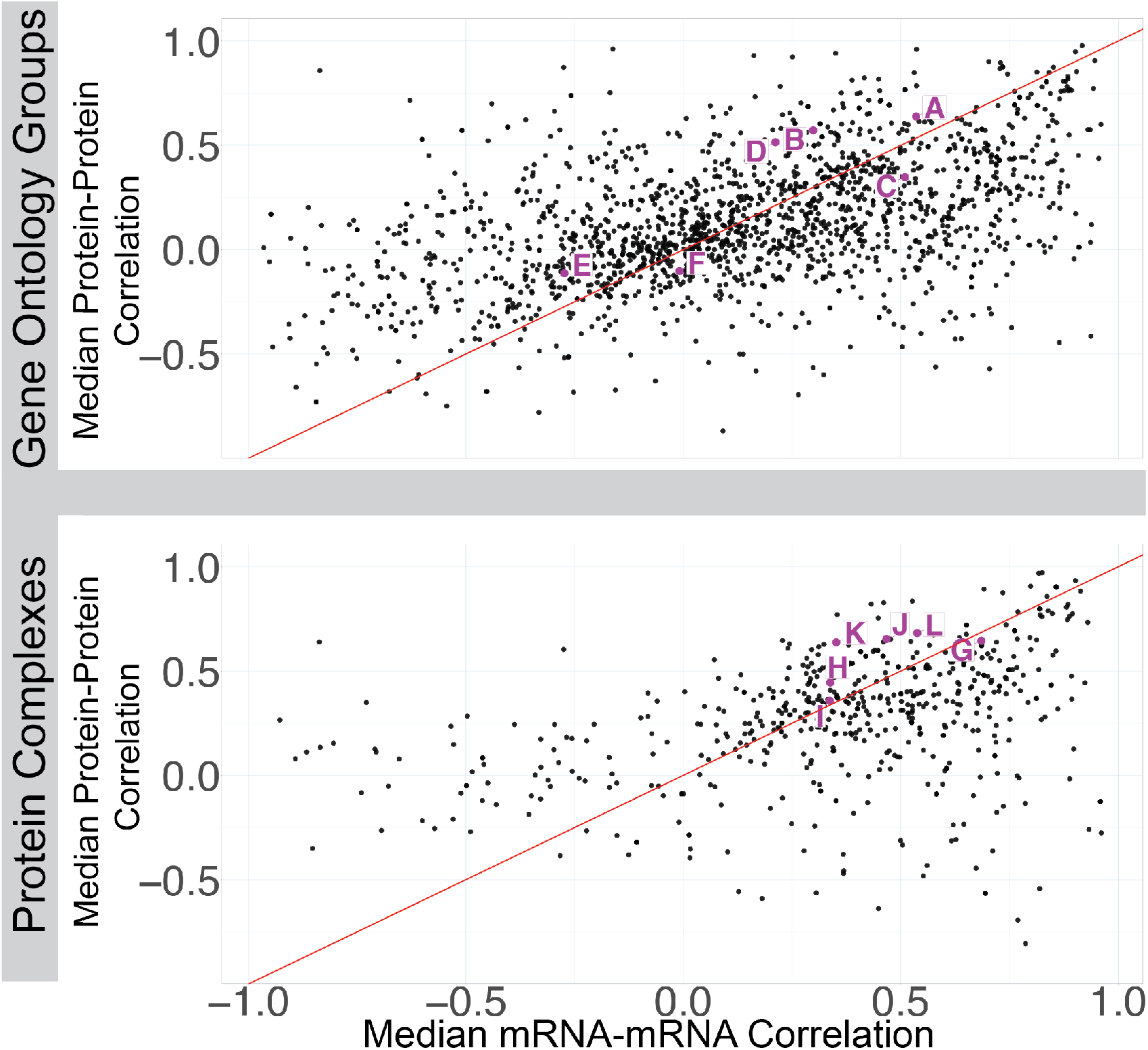
Comparison of pairwise mRNA-mRNA and Protein-Protein correlations for groups tested in Fig. 5. Protein-Protein correlations are the same as those in Fig. 5A. mRNA-mRNA correlations are computed in a similar fashion, where across clusters correlations are computed for all pairs of consensus transcripts in each group, and the median is computed across gene product pairs.

## Supplementary Data Tables

Details for data tables that were used to make the main section figures in the manuscript. The full set of all supplementary data is available here.

**Supplementary Table 1**: gene res.RData

Results from gene product level testing for significant rPTR. Data from the table was used for Fig. 2C. The table contains summary statistics for gene product level protein, mRNA and rPTR distributions generated via *BayesPG*.

**Supplementary Table 2**: filtered go rptr test.RData

Results from testing gene products which map to GO term for significant rPTR. Data from the table was used for Fig. 3 and Fig. 4. The table contains summary statistics for GO level protein, mRNA and rPTR distributions that were generated via *BayesPG*.

**Supplementary Table 3**: filtered complex rptr test.RData

Results from testing gene products mapping to protein complexes for significant rPTR. Data from the table was used for Fig. 3. The table contains summary statistics for Complex level protein, mRNA and rPTR distributions that were generated via *BayesPG*.

**Supplementary Table 4**: filtered go corr test.RData

Results from testing gene products which map to GO term for significant correlations across cell types. Data from the table was used for Fig. 5.

**Supplementary Table 5**: filtered complex corr test.RData

Results from testing gene products mapping to protein complexes for significant correlations across cell types. Data from the table was used for Fig. 5.

**Supplementary Table 6**: within go correlations summary.RData

Summary (mean and median) of the consensus protein-protein correlations of gene products which map to individual GO terms. Data from the table was used for Fig. 5.

**Supplementary Table 7**: within complex correlations summary.RData

Summary (mean and median) of the consensus protein-protein correlations of gene products which map to individual protein complexes. Data from the table was used for Fig. 5.

**Supplementary Table 8**: npop1 varMod ev updated.txt

Peptide x cell matrix that was processed the using variable modification search outputs. The data in this matrix was used to generate Fig. 6.

